# Sleep leads to system-wide neural changes independent of allo- and egocentric spatial training in humans and rats

**DOI:** 10.1101/2020.12.19.423580

**Authors:** Anumita Samanta, Laurens S van Rongen, Janine I. Rossato, Justin Jacobse, Robby Schoenfeld, Lisa Genzel

## Abstract

Sleep is important for memory consolidation, especially the process of systems consolidation should occur during sleep. While a significant amount of research has been done in regards to the effect of sleep on behavior and certain mechanisms during sleep, until now evidence is lacking that sleep leads to consolidation across the system. Here, we investigated the role of sleep in consolidation of spatial memory in the watermaze in both rats and humans using allocentric and egocentric based training. Combining behavior with immediate early gene expression analysis in rodents and functional MR imaging in humans, elucidated similar behavioral and neural effects in both species. Rats and humans showed a benefit of sleep on behavior. Interestingly, sleep led to systems-wide retrieval network in both species in both training conditions. Thus, we provide cross-species evidence for memory consolidation on the system-level occurring during sleep.

**Significance Statement:** Processes occurring during sleep such as memory reactivations are proposed to lead to consolidation from the initial hippocampal memory representation to long-lasting cortical representations, this is known as systems consolidation. By combining behavioral measurements in the watermaze with immediate early gene expression analysis in rats and function magnetic resonance imaging in humans, we could show a benefit of sleep on behavioral memory performance. And, sleep lead to systems-wide changes in the retrieval network. These results are the first direct evidence supporting the role of sleep for systems-wide memory consolidation in both rats and humans.

## Introduction

The ability to reliably navigate to known, desired locations requires integrating spatial information from different reference frames followed by consolidation of the information to build long-term spatial maps of the surrounding environment. Sleep has been proposed to have a special role in this later consolidation period [1-3]. Two memory systems are thought to be used to locate a target in space: a place learning (allocentric) and response learning (egocentric) system. Allocentric navigation, which relies on the development of a spatial cognitive map containing an internal representation of relations among distal cues, is known to be dependent on the hippocampus [4]. In contrast the egocentric system, which relies on the location of the navigator and may involve repeated use of relatively fixed motor movements to locate the target, is known to be dependent on the striatum [5]. During real world navigation, information from both frames are integrated to form a cohesive representation of the environment and the position of the navigator within [6]. However, one can bias the use of one strategy over the other by adopting specific training paradigms [7, 8]. Usually, this is done by including variable versus stationary starting locations within a maze, that will then bias to allo- vs. egocentric strategies respectively. Using allo- and egocentric training paradigms enables the investigation of specific initial memory circuits, hippocampus and striatum, and their associated consolidation processes.

Sleep is important for experiences to be consolidated to form long-term memories. It optimizes the consolidation of newly acquired information and has been proposed to reorganize brain circuits at both synaptic and systems level [1, 2]. Especially memory consolidation processes associated with the hippocampus, have been proposed to be dependent on sleep [9, 10]. On a systems level, the hippocampus is thought to be initially involved in the encoding of memories by binding different information stored in cortical modules into a coherent trace; and over time the connections of these cortical modules strengthen to thus become hippocampal independent [11, 12]. One critical mechanism underlying this process is thought to be repeated memory reactivations during NonREM sleep, which then lead to progressive strengthening of the cortico-cortical connections and thus consolidation across the system [1-3, 13]. These memory reactivations during sleep occur mainly during the hippocampal sharp wave ripple [3] and may be the reason why the hippocampus plays a special role in sleep-related memory consolidation [9, 10]. Considering the position of hippocampus as a crucial hub for spatial navigation and offline consolidation processes, it could thus be proposed that in a spatial context, allocentric learning would benefit more from sleep compared to egocentric learning. More importantly, while the role of sleep in consolidation across the system has been proposed since Marr [14] until now there is no direct evidence for this. Only indirect evidence has been provided, by comparing recently encoded memories (i.e. 30-60min) to more remote memories (i.e. 24h) with the difference hinting at a role of sleep [15]. However, this is then still confounded by the passage of time and not only the occurrence of sleep.

In this study, we aimed at adopting a translational approach to study the differential effects of sleep on allo- and egocentric memory representations in rats and humans. We used the watermaze [16], which has been a well-established paradigm to study different aspects of spatial navigation, especially contrasting allo- and egocentric training [7, 17, 18]. Further, a human analog of the maze has been developed and showed comparable performance in behavior across both species [19, 20]. Using the watermaze we tested for the influence of sleep on allo- and egocentric spatial memory training in both rats and humans and investigated the underlying neural signatures with immediate early gene expression analysis in rats and functional MRI in humans. We hypothesized an improvement in memory performance after sleep especially when trained under allocentric condition and that sleep would lead to a systems-wide consolidation process. We could confirm the expected behavioral effect of sleep with both rats and humans showing a benefit of sleep on behavioral performance that seemed larger after allocentric training. Interestingly, the neural effects did not show this specific effect of learning strategy. In both rats and humans, sleep but not wake led to systems-wide changes in the memory network after both allo- and egocentric training.

## Results

### Memory performance in rats and humans

Both rats and humans were trained with a one session paradigm in the watermaze to test for spatial memory [21]. For the rats the task consisted of eight training trials and a probe trial 20 hours later without the presence of a platform to test for long-term memory performance. They were divided into two training groups – allocentric and egocentric – with the main difference that the former started each trial from a different position while the latter always from the same point in the maze (Fig. 1A). After being trained under either of the conditions, they were further divided into sleep group (allowed to sleep in assigned sleep cages) and sleep deprived group (sleep deprived in their home cages for six hours after training by gentle handling, Fig. 1B) [21]. To assess memory performance, time spent in the target zone in relation to total time during the test trial was used. Analogous to the paradigm in rats, a virtual watermaze environment was used for the humans [19]. The environment setting consisted of two islands – cued and hidden – to enable subsequent functional MRI (fMRI) analysis. The cued island was a brown island with no distal landmarks and contained only a visible flag (cue) next to a treasure box (Fig. S1). The location of the flag kept changing every trial and the subjects were instructed to scan the area to find the target. The hidden island was a green island surrounded by four landmarks and a hidden treasure box, which was the target location (analogous to the platform in the watermaze). The box was hidden in a fixed location in a small indentation on the virtual island surface such that it would only be visible to the participants when they were close to it. The overall task was run as a block design with eight alternating trials of cued and hidden island resulting in a total of 16 blocks, that allowed us to isolate memory specific effects excluding for general visual input and movement through the virtual world in the subsequent fMRI analysis. Each trial was self-paced and ended with the participant marking the target location. The participants were allowed to freely navigate both islands with a joystick and their objective was to find the treasure box in each one and press a button on the joystick when they were in close proximity to the box. For the encounter with the hidden island, the participants were randomly allotted to either of the training conditions – allocentric or egocentric with changing or same starting position – and had to learn the location of the hidden box over the trials (Fig. 1A). This training and later test sessions was conducted in the MRI scanner. After the session, they were further divided into the sleep (take a nap with polysomnography for up to two hours (83.17±3.3 min, range 33.5-113.5 min), for sleep stage analysis see Fig. S2) or wake group. During the wake period the subjects were allowed to watch a neutral, non-emotional movie with an experimenter in the same room in order to monitor that the subject stays awake throughout the entire period. Following the sleep/wake intervention, the participants were taken back to the scanner and tested in the watermaze environment (Fig. 1B). In contrast to the rats for which the probe trial consisted of a single trial, the participants had to run all eight trials again in each island to enable the correct contrast in the fMRI analysis. However, in this session, they had to mark the location of the treasure box in the hidden island to the best of their knowledge without the box being present in every trial. In humans, the memory performance was measured as latency to reach the location. At test rats performed above chance across both sleep and sleep deprived groups for both allocentric and egocentric training conditions (Fig 1C, left panel). However, rats there was a general effect of sleep and an interaction of sleep and training condition on performance (sleep/sleep deprivation F_1,39_= 4.6, p = 0.039, allo/ego F_1,39_= 1.4, p = 0.244, interaction F_1,39_ = 3.8, p = 0.058). The human subjects were generally better in the egocentric in comparison to the allocentric condition, and as in rats showed a general effect of sleep on performance (allo/ego F_1,69_= 17.2, p < 0.001, sleep/wake F_1,69_ = 3.8, p = 0.056 and interaction F_1,69_ = 1.6, p = 0.2). In rats, the latency to reach the platform position at test showed a similar pattern as the dwell time analysis and human latency results, however it did not reach significance (Fig. S3, all p>0.2). In sum, in both rats and humans a sleep effect on behavior was seen, which was in rats more pronounced after allocentric training.

**Figure 1.**
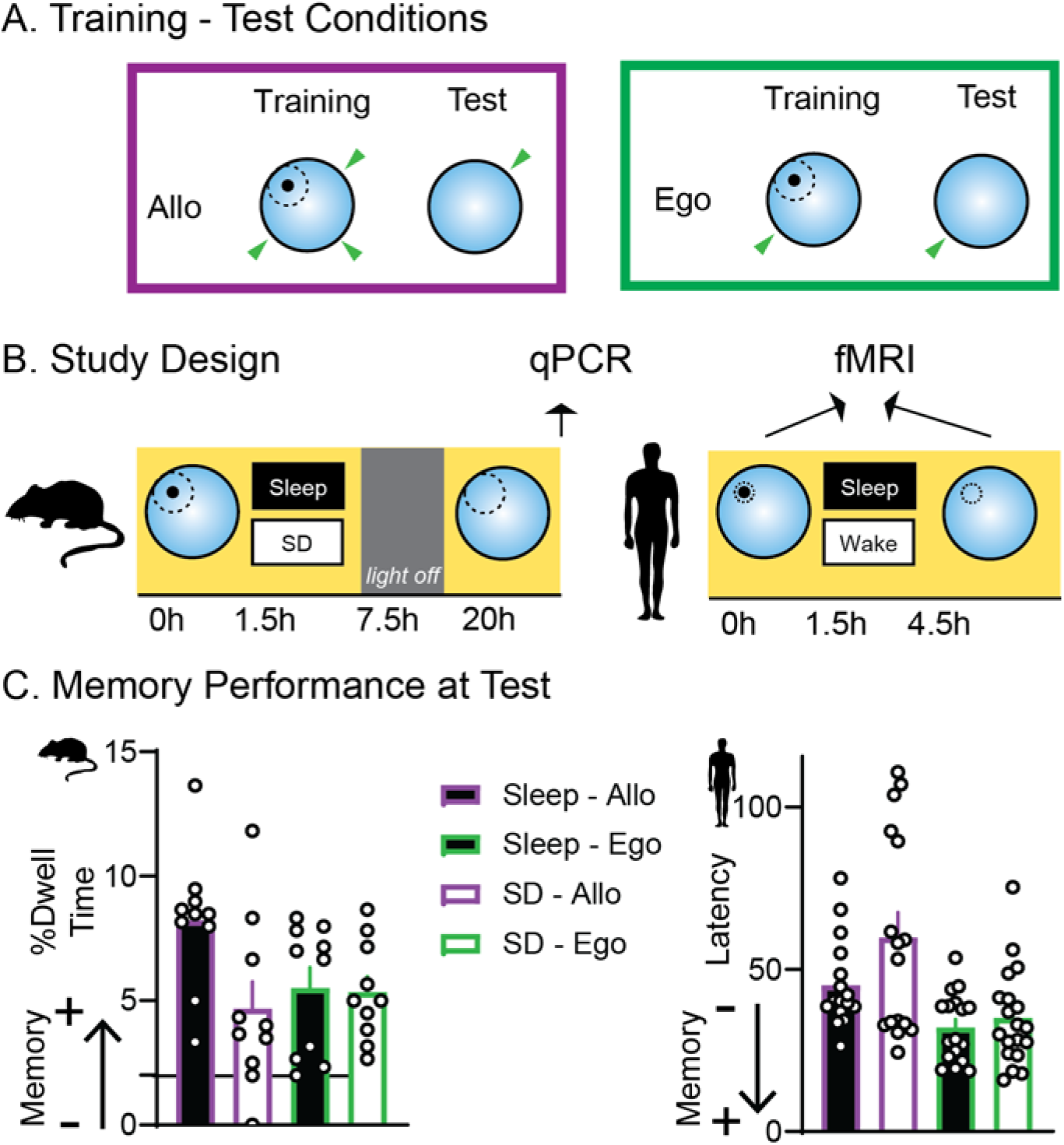
Study Design and Memory Performance. **(A)** The training-test conditions. The left panel shows the allocentric condition during which both rats and humans started from different locations on every training trial. In contrast, during the egocentric condition the starting location remained the same for every training trial (right panel). **(B)** Shows the study design for rats (left) and humans (right). Rats and humans were divided into two groups based on the training condition (allo/ego) and underwent a single day training protocol in the watermaze with eight training trials. After training, each group was subdivided into two additional groups: sleep (allowed to sleep individually in sleep cages with video monitoring) and sleep deprived (SD) group (sleep deprived for six hours after training by gentle handling in the home cage) in rats and sleep (took a nap with polysomnography for up to two hours, for sleep stages see Fig. S2) and wake (watched a neutral movie for two hours) in humans. Overall, there were hence four groups each of rats and humans: sleep-allocentric (n=10), sleep-egocentric (n=10), SD-allocentric (n=10), SD-egocentric (n=10) in rats and sleep-allocentric (n=17), sleep-egocentric (n=16), wake-allocentric (n=16), wake-egocentric (n=19) in humans. Sleep/SD in rats ended when the light-on period switched to the light-off period. Rats from all groups were tested the next day at the onset of the light-on period with a probe trial in the watermaze (no platform present). 30 min after the test trial, the rats were killed and brain regions were collected (prefrontal cortex, hippocampus and striatum) for qPCR analyses on retrieval induced expression of different immediate early genes. Humans from all groups were tested after the sleep/wake session in the watermaze environment with fMRI. **(C)** Behavior results of the rats (left) and humans (right). All rats performed above chance level (chance level is 2%) with those animals that slept after training showing a significant improvement in performance in contrast to those awake in dwell times (higher values indicate better memory). The human subjects, who took a nap between training and test, showed a significant improvement in memory performance compared to those that stayed awake (lower values indicate better memory). The purple bar contours used for the allocentric condition and the green contours for the egocentric condition. The black and white filled bars for both colors correspond to sleep and sleep deprived (SD) group respectively. Error bars are SEM.

### Retrieval induced IEG expression analyses in rats

After establishing the behavioral effect of sleep, we next moved on to test the neural effects. For this, in rodents, we measured the retrieval-induced expression of immediate early genes. More specifically we measure expression of *Arc, cFos* and *Zif268* in the prefrontal cortex, striatum and hippocampus. Immediate early genes expression can be used as an index for neuronal activation [21-24]. In a full model including gene and brain area as within subject factors and sleep and training type as between subject factors, there was a significant effect of sleep and gene and an interaction between training type (allo/ego) and brain area, gene and sleep as well as gene and brain area (sleep F_1,16_ = 8.5, p = 0.01; training type X brain area F_2,33_ = 4.1, p = 0.026; gene F_2,32_ = 4.7, p = 0.016; gene X sleep F_2,32_ = 3.1, p = 0.061; gene X brain area F_2.6,42.3_ = 3.7, p = 0.023; other F<1.9 p>0.13; Fig. 2). An interesting pattern emerged in which sleep led to an increase in gene expression in all brain areas (prefrontal cortex, hippocampus and striatum) whereas after sleep deprivation animals only showed increased gene expressions in the hippocampus for allocentric and striatum for egocentric training. Thus, only after sleep systems-wide memory retrieval could be seen, which was independent of allo- or egocentric training conditions. In contrast, if animals were sleep deprived after training, memory retrieval was associated only with those brain areas that are known to be necessary for each training type: striatum for egocentric and hippocampus for allocentric.

**Figure 2.**
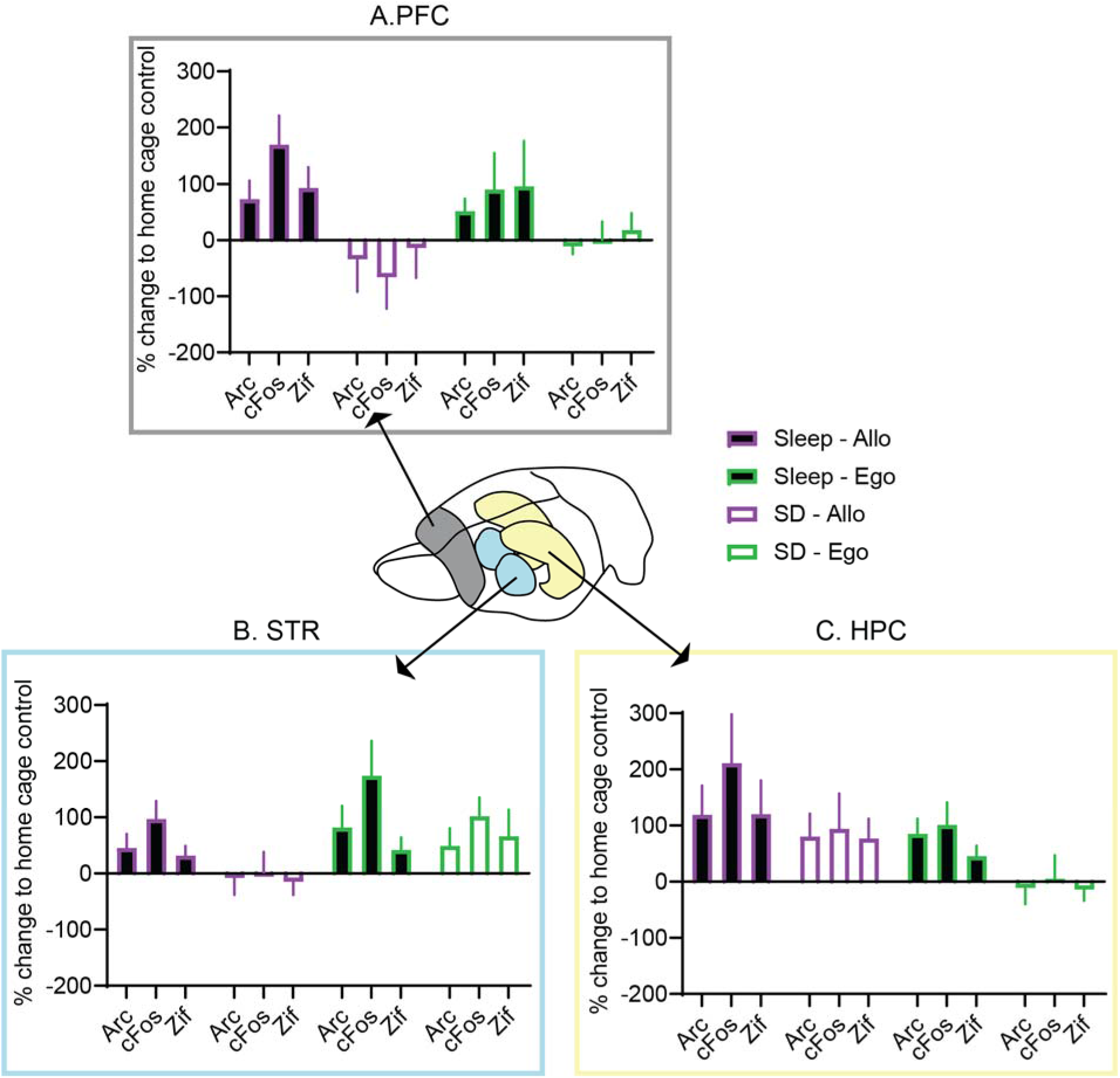
Memory Retrieval in Rats. Here, the gene expression profile for immediate early genes *Arc, cFos* and *Zif268* (represented as % change in relation to home cage controls) across different brain regions for all groups of rats. **(A)** Expression profile in prefrontal cortex (PFC), significant more gene expression of all genes is seen for the sleep groups in contrast to the sleep deprived (SD) groups. **(B)** In the striatum (STR) both sleep groups as well as egocentric sleep deprivation group showed increased gene expression in comparison to sleep deprivation allocentric. **(C)**. In the hippocampus (HPC) both sleep groups and sleep deprivation allocentric showed increased gene expression in comparison to sleep deprivation egocentric. The purple bar contours used for the allocentric condition and the green contours for the egocentric condition. The black and white filled bars for both colors correspond to sleep and sleep deprived (SD) group respectively. Error bars are SEM.

### Effect of sleep on brain activity in humans

Next, to assess the neural correlates of sleep on spatial memory under allocentric and egocentric training in humans, we analyzed the MRI BOLD images acquired during the training and test session. In both sessions, subjects completed eight training trials to a fixed treasure in the hidden island under either allocentric or egocentric condition as well as eight trials to navigate to a visible flag in the cued island, which position changed from one trial to the next. Thus, the first-level contrast was between the hidden and cued island to enable isolation of memory encoding and retrieval specific effects while controlling for the general task properties such as joystick movement and visual input. Only the first 30s of each trial were included in the analysis, to control for the fact that due to self-pacing of the trial each trial length was different and there was a difference in average latency at test over the groups (mean 42.5 s; range 15.8-110.6 s). However, including the whole trial periods did not affect the general results (see Fig. S6-8). Only when subjects slept between training and test, significant changes were seen in BOLD activity with increases in medial and lateral frontal cortices, anterior and posterior parietal cortices, visual cortex, cerebellum and a decrease in activity in medial prefrontal cortex, precuneus and hippocampus (Fig. 3 shows the contrast between training and test for sleep both training groups, for sleep allocentric and sleep egocentric separately see Fig. S4, 5, 7, 8). All results were collected at uncorrected p<0.005 and then corrected on the cluster level to control for multiple comparisons with p<0.05 FWE (full-factorial model with factors training-test, allo-ego and sleep-wake). It is noticeable that brain areas that showed an increase due to sleep belong to the executive control network, which has previously been shown to be related to goal directed behavior [25] and spatial memory [26]. In contrast, the brain areas that showed a decrease belong to the default mode network, which has also been previously associated to spatial memory [27-31].

**Figure 3.**
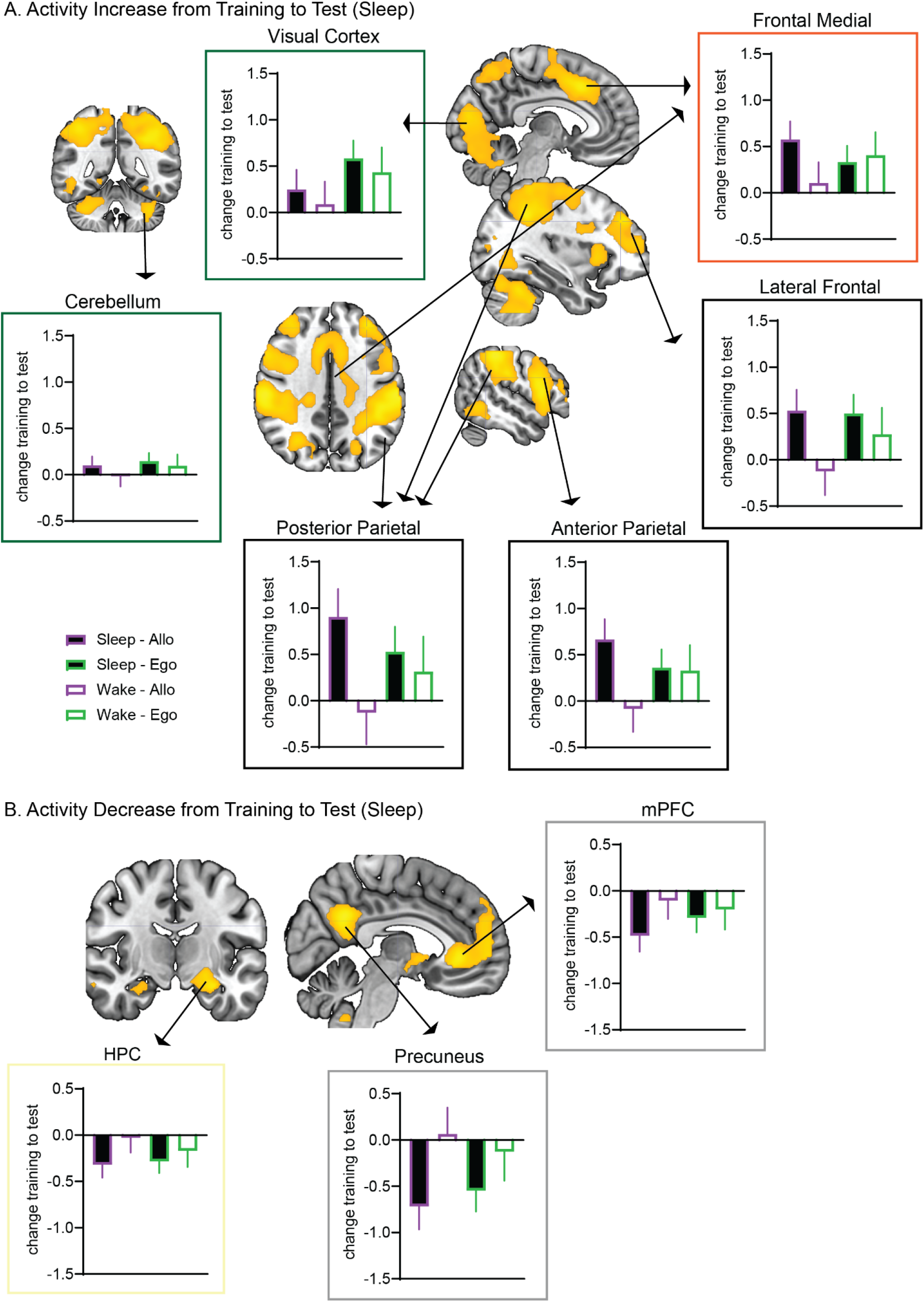
fMRI Activity Analysis in Humans. **(A)** Brain maps show the changes from both sleep groups (test more than training, p<0.05 FWE-cluster corrected) and bar graphs are extracted for each subgroup for the peak voxel of each cluster. An increase in activity was observed in the medial and lateral frontal cortex, anterior and posterior parietal cortex. Increases were also observed in a part of the visual cortex and cerebellum. This activity increase was similar across both training conditions over sleep (maps for each group see Fig. S4, 5, 7, 8). In contrast, after wake the whole brain analysis maps were empty. However, the peak voxel activity extraction indicates that egocentric wake did show similar changes to sleep, even if the smaller change and larger variability in this group precluded significance on the whole brain level. **(B) B**rain maps show the changes from both sleep groups (test less than training, p<0.05 FWE-cluster corrected) and bar graphs are extracted for each subgroup for the peak voxel of each cluster. A decrease in activity was observed in the medial prefrontal cortex (mPFC), precuneus and hippocampus (HPC). The purple bar contours used for the allocentric condition and the green contours for the egocentric condition. The black and white filled bars for both colors correspond to sleep and sleep deprived (SD) group respectively. Error bars are 95% confidence interval. For detailed statics on each cluster, see Tables S1-2.

The same contrasts for the wake subjects were empty (both for each training group separately as well as for the combined wake group). However, when signal change at the peak voxel for each cluster was extracted for each group separately, similar changes as shown in the sleep groups could be seen for the wake egocentric group, even if they were not significant in whole brain analysis (also not seen uncorrected p<0.005). Additional contrast were run to further support of the difference between allo- and egocentric changes across wake. At test, only for allocentric and not for egocentric, the same areas that showed increases from training to test during sleep, were also more active in subjects that slept versus those that stayed awake (allo sleep > allo wake at test, Fig. S9). Further, the interaction of sleep versus wake and training to test changes showed no significant results when including both ego- and allocentric or if only ego subjects were included. However, this same interaction (sleep/wake and training/test) was significant if only the allocentric subjects were included, with areas belonging to the executive control network showing increases in sleep but not wake (allo sleep training < allo sleep test and allo wake training > allo wake test, Fig. S10).

In summary, similar to the findings in rats, we see a change in the whole brain independent of training after sleep, which is not present after wake. More specifically, we observed a shift in brain activation after sleep with an increase in in regions belonging the executive control network and a decrease in regions belonging to the default mode network also including the hippocampus. Similar changes could be seen in egocentric wake group, but these were much weaker and did not reach significance. Further, these changes were absent in the allocentric wake group.

### Connectivity changes in humans

The brain areas that showed significant changes in the activity analysis are known to be part of brain networks with areas showing increases belonging to the executive control network and areas showing decreases associated with the default mode network. Thus, we next investigated functional connectivity changes during task execution by conducting a Psychophysiological Interaction (PPI) analyses. Led by the activity analysis we focused on the medial frontal cortex as region of interest (ROI −2 −10 48) for this analysis. This ROI is a key hub of the executive control network, related to goal directed behavior [32]. As with the activity analysis, we only included the first 30s of each trial and all results were collected at uncorrected p<0.005 and then corrected on the cluster level to control for multiple comparisons with p<0.05 FWE (full-factorial model with factors training-test, allo-ego and sleep-wake). At test in contrast to training the medial frontal cortex was functionally connected less to areas known to be part of in the default mode network but again only for those subjects that slept and not those that stayed awake (inferior parietal cortex, precuneus, prefrontal cortex, hippocampus, Fig. 4). Further, for both sleep and wake the cerebellum decoupled from the medial frontal cortex (for wake contrast see Fig. S11). As with the activity analysis, we extracted the change in connectivity for the peak voxel in each cluster for each group separately.

**Figure 4.**
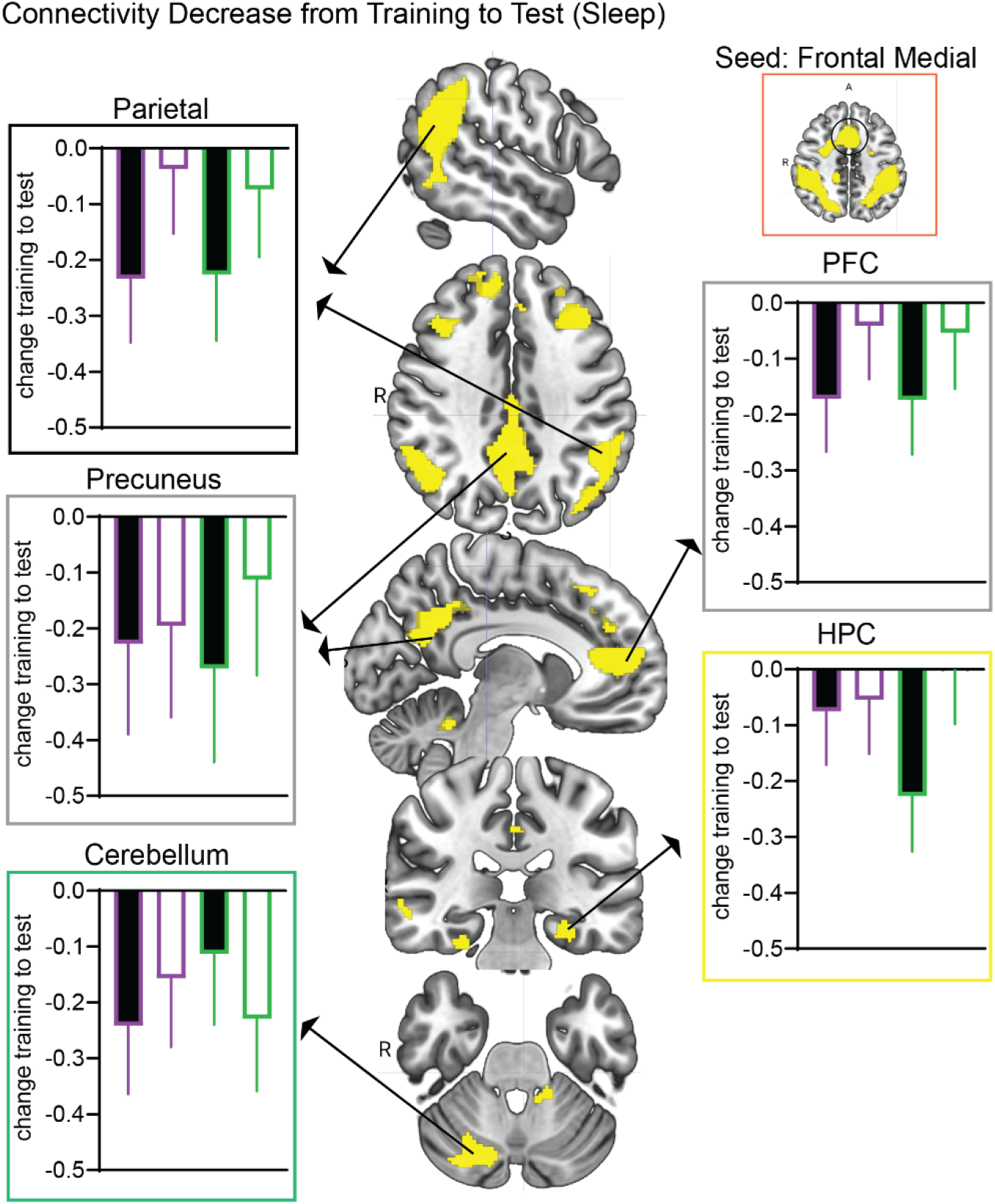
fMRI Connectivity Analysis (PPI) in Humans with Medial Frontal ROI. **B**rain maps show the changes from both sleep groups (test less than training, p<0.05 FWE-cluster corrected) and bar graphs are extracted for each subgroup for the peak voxel of each cluster The frontal medial cortex showed a significant decrease in connectivity with the prefrontal cortex (PFC), hippocampus (HPC), precuneus and inferior parietal cortex from training to test (see Table S3). These regions are known to be part of the default mode network. Additionally, it also a significant decrease in connectivity with the cerebellum for both sleep and wake groups (wake contrast in Fig. S11). The purple bar contours used for the allocentric condition and the green contours for the egocentric condition. The black and white filled bars for both colors correspond to sleep and sleep deprived (SD) group respectively. Error bars are 95% confidence interval.

The decreased connectivity and thus decoupling between the main hub of the executive network and default mode network, parallels the finding in the activity analysis with increase seen in the former and decreases seen in the latter network across sleep but not wake period.

## Discussion

In this study, we investigated the effects of sleep and wake on allocentric and egocentric memory representations in rats and humans using the watermaze. Overall, memory was intact independent of training condition or if subject slept or stayed awake between training and test. However, in both rats and humans sleep led to better memory performance in comparison sleep deprivation. This effect of sleep seemed larger in the allocentric training group, however the interaction between sleep and condition was only significant in rats. Human subjects in the egocentric condition were in general better than those in the allocentric training condition. To investigate effects on brain activity two different methods were used in the different species: in rodents, we measured retrieval-induced immediate early gene expression in the hippocampus, striatum and prefrontal cortex in comparison to home cage control. In contrast, in humans we compared the MRI BOLD signal at training and test. These analysis showed that in rats and humans a systems-wide change was seen after sleep for both allo- and egocentric training conditions, which was not present after wake.

More specifically in rats, after sleep increases in retrieval-induced gene expression was seen in all three brain areas (hippocampus, striatum, prefrontal cortex), which was independent of allo- or egocentric training conditions. In contrast, when the rats were sleep deprived after training, memory retrieval was associated with increased gene expression only in those brain areas that are known to be necessary for each training type: striatum for egocentric and hippocampus for allocentric. In humans, fMRI analyses only showed significant changes from training to test in subjects that slept and not those that stayed awake, with significant increases in activation in areas that are associated with the executive control network (such as superior posterior parietal cortices, frontal medial cortex), and significant decreases in areas associated with the default mode network (hippocampus, medial prefrontal cortex, precuneus). Extracting activity changes for the peak voxel in each cluster for each group revealed that the egocentric wake condition showed similar changes as both sleep groups, however these changes were not visible or significant in the whole brain analysis (also uncorrected p<0.005). Further, only for allocentric was there a significant difference between wake and sleep at test as well as significant interaction between wake/ sleep and training/test. Connectivity analyses during the task with PPI showed a functional decoupling of the frontal medial cortex – the key hub of the executive control network – with areas of the default mode network. Thus, we observed across both species systems-wide changes across sleep that were the same for both allo- and egocentric training, in contrast wake had differential effects on both training conditions.

Sleep is crucial for offline consolidation processes and strengthening memories. The proposed key underlying mechanism is neural reactivation wherein neural activity present during encoding reemerges during NonREM sleep [2, 3]. These reactivation events have been shown to occur mainly during hippocampal high frequency burst oscillations, referred to as sharp wave ripples [3, 33]. Further, these offline consolidation processes are thought to involve a dialog between the hippocampus and cortex via the hippocampal sharp wave ripple in combination with the neocortical slow oscillation and sleep spindle, which should stabilize the labile memory traces in the cortex leading to systems consolidation [1, 3, 8, 12, 13, 34, 35]. Considering the crucial role of hippocampus in spatial navigation and allocentric learning [36], it has thus been proposed that memories encoded with allocentric training – known to depend on the hippocampus – would benefit more from sleep [9, 10]. In contrast, egocentric learning is dependent on the striatum and thus consolidation of these memories should be less dependent on sleep [5, 8]. Several studies did provide evidence for this dissociation [37-39]. Hagewoud and colleagues [39] sleep deprived rodents after learning a plusmaze and could show a shift to an egocentric strategy from a preferred allocentric learning strategy in animals that could sleep; this was accompanied by a respective shift from hippocampal to striatal levels of retrieval-induced pCREB. In humans, the allo- vs egocentric strategies has only been investigated using a motor-sequence task, during which the use of respective strategy can be tested if the finger tapping sequence is expressed in the other hand corresponding to global features (allocentric, left ring to index finger switch would be right index to ring finger switch) or internal features (egocentric, left ring to index finger switch would be right ring to index finger switch). These studies could show an improvement in performance for the allocentric strategy, in contrast to the egocentric memory expression, which was maintained over wake [37, 38]. A similar effect of sleep on memory performance according to training strategy, could be seen in our rats as well as humans. All subjects trained under allocentric condition show an improvement in performance over sleep, and the sleep-effects on behavior were numerically less for rats and humans trained under egocentric condition. However, only in rats was the interaction between training type and sleep significant. In contrast, in humans the subjects in the egocentric condition performed generally better. For both rats and human a main effect of sleep on performance was significant.

Interestingly, the neural effects showed a different pattern. In both rats and humans a systems-wide change was seen across sleep but not wake, which was the same for both training conditions. Thus, while theory proposed that only hippocampal dependent memories would benefit from sleep, perhaps the different memory systems may be more interrelated than previously thought, especially during sleep. The ventral striatum has been proposed to integrate inputs from the hippocampus, prefrontal cortex and related subcortical structures to construct outcome predictions and stimulate goal directed behaviors [40, 41]. And, while memory reactivations are most known for the hippocampal systems, Pennartz and colleagues have shown memory reactivations in the striatum, which were in close temporal association with hippocampal ripples [40, 42, 43]. Thus, perhaps it is less surprising that we see systems-wide neural changes after sleep in both allo- and egocentric training conditions. These findings fit well with the proposed role of sleep for systems-wide consolidation and thus perhaps consolidation of memories independent of learning strategy to allow for future flexible use and adaptability [1, 3, 8, 13]. Further, this data is the first of its kind to support the role of sleep for systems level consolidation while controlling for the general passage of time.

Our fMRI results show a dynamic interaction between areas of the executive control network and the default mode network over sleep across both training conditions. We observe an increase in areas belonging to the executive control network and a decrease over sleep in areas belonging to the default mode network that includes the hippocampus. The executive control network has been shown to be active during attention demanding visuo-spatial tasks, goal directed behaviors and navigation with the parietal cortex especially implicated to play a critical role in sensorimotor integration and activities of higher cognitive function [44]. The posterior parietal cortex is of particular interest with respect to spatial navigation tasks and has been shown to be the region coding for sensory locations of stimuli into appropriate motor coordinates required for making directed movements [6, 32]. Multiple studies in primate and rat models have also indicated the role of posterior parietal cortex in encoding route progression during navigation under both allocentric and egocentric conditions, adapting to external environments and maintaining an internal cognitive map of self-position in space [45, 46], also when tested in virtual environments [47]. The posterior parietal cortex serves as a cortical integration site for hippocampally generated allocentric spatial information and egocentric spatial orientation to permit goal-directed navigation [48-51]. Further, memory reactivations during sleep have been observed in the parietal cortex as in the prefrontal cortex [52, 53] and both brain areas show high frequency oscillations during NonREM sleep cooccurring with hippocampal ripples [51]. Consistent with these findings, we observe an increase in activation in posterior parietal cortices across both training conditions after sleep.

In regards to the changes seen in areas of the default mode network, ample amount of evidence has pointed to the importance of hippocampus and medial temporal lobe structures including the prefrontal cortex and precuneus (also known as retrosplenial cortex [54]) in spatial navigation in both human and rodent models [26, 55-57]. Furthermore, the default mode network is functionally modulated by sleep with persistent connectivity during light sleep and especially sleep spindles and gradually decoupling with deep sleep [34, 58-61] and thus could potentially play a role in offline consolidation processes coordinating the systems-wide consolidation process during sleep [8, 29-31, 62]. Recent evidence in rodents show the co-occurrence of cortical high-frequency oscillations in default mode network areas as well as posterior parietal cortex with hippocampal ripples during NonREM sleep [51], which may be the mechanisms how memories could be consolidated from the initial hippocampal storage to downstream areas such as the posterior parietal cortex via the cortical default mode network areas [8]. This could be the mechanisms underlying our findings, where we see a shift in activity with decreases of the default mode network, which is perhaps the intermediate storage, and increases in the goal directed network including the parietal cortex over sleep.

With these results, it is important to consider the source of the neural activation measure and baseline contrast for each species. For the rats, we measured retrieval induced changes in gene expression of immediate early genes in the hippocampus, striatum and prefrontal cortex in contrast to home cage control. Immediate early genes show increased expression in those cells that are especially active at a given moment and can be used to test for activity related to memory retrieval [21]. The results in rats thus highlight which brain areas are more active during memory retrieval in comparison to behaviors in home cage controls. For the humans, we measured BOLD responses both during encoding and retrieval of the task and the results focus on relative changes in regions active during each of the sessions. The prime measure however is relative since we first create contrasts of regions active during hidden versus cued island and then on the second level changes from first session to the second one. Further, the BOLD signal, while also known to measure brain activity, is based on blood oxygenation changes and is thus a more indirect measure than gene expression used in rats. The main difference in results in both species would be that for rats, the expression levels are compared to those in home cages whereas for the humans, the comparison is memory specific (cued vs hidden island) and between sessions (within subject). Keeping the these limitations in mind, we thus here in the discussion rather focused on a rough comparison of differences seen across training type and sleep groups and had less focus on directly comparing differences between the species.

In sum, across both species and training conditions we see systems-wide retrieval network after sleep which is not present after wake fitting to the main significant effect of sleep on behavior, even though behavior effects of sleep was more pronounced in the allocentric condition. Thus, we are the first to provide cross-species evidence for the proposed function of sleep for systems-wide consolidation of memories already proposed by Marr [14].

## Acknowledgments

We would like to thank Elise Marie Solsnes, Kellie Moffat, Chloe Stephenson-Wright, Antonis Asiminas, Adrian Duszkiewicz, Tomonori Takeuchi, Elisabeth Allison and Roddy Grieves, who helped with the rodent experiments in Edinburgh and Svenja Rohde for the sleep scoring of the human nap data. We would also like to thank Guillen Fernandez, Niels Kohn, Ruud Berkers, Paul Gaalman, Jessica Askamp and Marcel Zwiers for help with the MRI and human EEG data acquisition and analysis. The study was funded by the Branco Weiss Fellowship – Society in Science to Lisa Genzel.

## Contributions

A.S. performed the human experiments and wrote the first draft of the manuscript., L.v.R. helped with the data analysis of the fMRI data, J.R. and J.J. performed the qPCR analysis in the rodents, R.S. developed and adapted the human testing environment, L.G. designed the project, supervised all experiments and analysis and co-wrote the manuscript, all authors contributed to the final revisions of the manuscript.

## Methods and Materials

### Subjects

#### Rats

Adult male Lister-hooded rats (Charles River, United Kingdom), aged 8-10 weeks at the start with an average weight of 250-300 g were used for the experiment. They were housed in groups of four animals per cage, with free access to food and water at all times, and kept on a delayed day-night cycle (10 A.M. – 10 P.M. light on). After arrival, the animals were habituated to the housing environment for at least a week and then handled across 3 days with at least 5 mins each day before starting with watermaze habituation. A total of 45 rats were used from which 25 were used for qPCR experiments. All experimental procedures were in accordance with national (Animals [Scientific Procedures] Act, 1986) and international (European Communities Council Directive of 24 November 1986[86/609/EEC]) legislation governing the maintenance of laboratory animals and their use in scientific experiments. The minimal number of rats for the necessary statistical power was used, with random assignment to groups, and minimal suffering was ensured for all the experimental procedures. The experiments were approved by the UK home office under the project license number 60/4566 and the Experimental Request Forms by the Edinburgh University division of the National Veterinary Service.

#### Humans

Seventy-seven neurologically healthy, right-handed male subjects (age range = 18-30 years, mean = 24) were recruited for the study. Only males were chosen since only male rats were used in the previous experiment. Additionally, sex differences studies across both rats and humans [20] have shown sex and menstrual cycle to influence sleep dependent memory consolidation and watermaze training. Participants were recruited through the Radboud Research Participation System. All provided written informed consent prior to the start of the experiment and were paid for their participation. This study was approved by the local ethics committee (CMO Arnhem-Nijmegen, Radboud University Medical Center) under the general ethics approval (“Imaging Human Cognition”, CMO 2014/288), and the experiment was conducted in compliance with these guidelines. Exclusion criteria for the participants were taking sleep medications, regular naps and involved in professional gaming activities. They were screened for these criteria before the start of the session. Additionally, their alertness levels and sleep quality were assessed for with the Stanford Sleepiness Scale and Pittsburgh Sleep Quality Index respectively during the experiment session. Values per group for these measures are tabulated in Table S1. Eight subjects were excluded from the experiment. For five of them, the joystick was incorrectly calibrated, and for the remaining, there were technical problems including abrupt crashing of the task environment program during the scan.

#### Watermaze Task - Rats

For both habituation and training, the procedures were adopted from [21]. Prior to the start of the main experiment session, the rats were first habituated with a visual-cue version of the watermaze (diameter = 2 m) for three days, with four trials per day. They had to find the submerged platform in the water maze, indicated by a visual cue placed on top of the platform (diameter = 12 cm), while curtains surrounding the pool hid any extramaze cues. On reaching the platform, the rats had to wait on the platform for 30 seconds before being picked up and then continued with the next trial. In the end, the rats were familiar with the procedure before beginning of training in the maze

A one session training design was adopted for this experiment as used in [21]. The session consisted of eight training trials followed by a probe trial after 20 hours. The rats were divided into two training groups – allocentric and egocentric. In the allocentric condition, the rats were placed in the watermaze from a different start quadrant each trial (facing the wall) and had to reorient themselves to locate the platform. In the egocentric condition, the rat had the same start location each trial to locate the platform. Platform locations and start positions were counterbalanced across animals. For both training conditions, if the rats did not reach the platform by 120 seconds, they were guided to the location. Distal extra maze cues were present to help the rats orient themselves in the water maze. After the session, the rats were randomly allocated to one of the two conditions sleep by allowing them to sleep in allotted sleep cages or sleep deprivation by gentle handling for 6 hours after training in their home cages. Gentle handling included handling the rats occasionally, gently tapping on the cage or removing the cover as soon as the animal started showing signs of tiredness [21]. 20 hours later, the memory performance of the rats were tested with a single probe trial, where they were placed for 120 seconds in the watermaze without any platform present and then picked up from the pool from the current location while their swim paths were tracked by an automated software (Watermaze, Watermaze Software, Edinburgh, UK [63]). The experiments were times so that sleep/sleep deprivation ended at light off (thus the transition to the active period) and the test was conducted at light on. Thus the animals had 12h to recover from the intervention before test, but sleep rebound was minimized by using the active period. After the test trial, the rats were sacrificed and brain regions (prefrontal cortex, striatum and hippocampus) were extracted for qPCR studies. Each 20 animals were trained in allocentric and egocentric conditions, with each half sleep or sleep deprivation, resulting in n=10 per subgroup. Of these 5 rats from each group were randomly selected as representatives for the qPCR analyses. In addition, there were 5 home cage rats who did not undergo behavior training and were used as home cage controls in the qPCR analysis.

#### RT qPCR Analysis - Rats

For qPCR measurements see also [21], the rats were sacrificed 30 min after the probe test trial. The home cage controls were also sacrificed at the same time point as the experimental rats. Immediately after extracting the brains, the medial prefrontal cortex, striatum and bilateral hippocampi were dissected and flash frozen with liquid nitrogen and stored at −80°C for later processing. Briefly, samples were homogenized and RNA was obtained via phenol-chloroform extraction according to the manufacturers’ instruction. Next, cDNA was synthesized in vitro with use of random hexamers. Subsequently, a RT-qPCR and comparative Ct quantitation was performed in experimental duplicates for *cFos, Arc, Zif268*, and 18S ribosomal RNA as the internal control on a StepOnePlus (Applied Biosystems, Carlsbad, United States) PCR machine. Plates were counterbalanced and amplification thresholds set manually (StepOne Software Version 2.3, Life Technologies). The amplified product size was verified using gel electrophoresis and amplification checked for primer-dimer formation and nonefficient DNase treatment. Data was normalized to the internal control 18S (also known as Rn18s, coding for ribosomal RNA47), and subsequently “fold” and then “percentage change” to home cage control or other control was calculated. Percent (%) change was used for statistical analysis and graphical presentation because fold change cannot be used for statistics and percent change gives a more intuitive sense of effect sizes.

#### Study Design - Humans

The entire experiment session lasted for a maximum of 6 hours and was split into three sub-sessions: 1, fMRI session with the subject being trained on the task; 2, 2.5-3 hour interval involving either taking a nap with EEG or watching a neutral movie; 3, followed by a second fMRI session where they got tested in the task. The session started at noon with the subjects filling in a couple of screening questionnaires and rating their alertness levels on the Stanford Sleepiness Scale. After having met with all inclusion criteria, they started with the first fMRI session. It started with a T1-weighted anatomical scan, followed by a Resting State scan where the participants were asked to fixate on a cross projected on the screen. Next, they performed 8 blocks of the training set (allocentric or egocentric alternating with cued island). The duration of this scan varied across all participants and ended when they successfully completed all the blocks. Following the task, the Resting State scan was repeated and then the fMRI session ended. At the end of the first fMRI session, the subjects were randomly allocated to either of the two conditions – wake (to watch a neutral movie for 2 hours with an experimenter present in the same room to monitor that the subject stays awake throughout the period) or sleep (take a 1.5-2-hour nap with Polysomnography). At the end of either of the movie/nap, participants were asked to rate their awareness levels again on the Stanford Sleepiness Scale and they started with the second fMRI session. As with the first round, this session started with a Resting State Scan followed by 8 blocks of test set (allocentric or egocentric alternating with cued island). The duration of this scan also varied across all participants and ended when they successfully completed all the blocks. Following the task, the Resting State scan was repeated and then the second fMRI session ended. This marked the end of the full experimental session. Participants from the sleep condition were asked to come another day for a second session where they had to take a short nap with Polysomnography (data not shown here).

#### Virtual Watermaze (VWM) Task - Humans

Analogous to the rats, humans were also trained in a watermaze environment to test for spatial abilities. For this purpose, we employed a virtual watermaze [19, 64], which consisted of a virtual island surrounded by four landmarks (distal cues) - a bridge, a sailboat, a wind turbine and a lighthouse (see Fig. S1). There was a hidden treasure box on the island, which was marked as the target location (equivalent to the platform in the water maze). The box was hidden in a fixed location in a small indentation on the virtual island surface such that it would only visible to the participants when they were close to it. This island would henceforth be referred to as the “hidden island”. The setup consisted of another island with no distal cues for orientation except for a visible colorful flag (cue) next to a treasure box, which position kept changing across each trial and was visible from a distance. This island would henceforth be referred to as the “cued island”. The overall task design was a block design with 8 alternating blocks of cued and hidden island resulting in a total of 16 trials. Each trial was self-paced and ended with the participant marking the target location. There was a 15 second interval between the finish of one trial and the start of another, during which the subject could turn around in the maze and orient themselves. The participants were allowed to freely navigate both islands with a joystick and their objective was to find the treasure box in each one and press a button on the joystick when they were in close proximity to the box. They would first encounter the cued island and had to find the visible flag. This island was used as a control to control for motor and visual input as well as isolate memory effects in the fMRI analysis. For the encounter with the hidden island, the participants were randomly allotted to either of the training conditions – allocentric or egocentric. The subjects were not aware of these two possible conditions. In the allocentric condition, they would have a different start location every trial and would have to reorient themselves each time to find the target location, thereby promoting place navigation. In the egocentric condition, they would have a fixed start location every trial, hence could rely on a repeated fixed movement to get to the target location in addition to using the visible cues. The main objective of the participants in both conditions was to learn the fixed location of the target box across all the trials. Finishing all 16 trials would mark the successful completion of the training set. For the test set, the island setup remained the same with one modification – the treasure box was removed from the hidden island and the participants were instructed to mark the location to the best of their knowledge, where they recalled the box to be located.

#### Polysomnographic Recordings

For the nap condition in the afternoon, polysomnographic recordings were obtained with 250 Hz sampling frequency, a 0.3 Hz high-pass filter, and a 35 Hz low-pass filter (BrainAmp, Brain Products, Gilching, Germany). 32 scalp electrodes were prepared including Fz, F3, F4, Cz, C3, C4, Pz, P3, P4, Oz, O1, O2 electrode sites and referenced to the left mastoid. Additionally, horizontal and vertical eye movements (EOG), electromyogram (EMG) on the chin and electrocardiogram (ECG) were recorded. Sleep scoring was performed by an experimenter blinded to the conditions, based on EOG, EMG and the following channels – F3, F4, C3, C4, O1, O2 using 30 second epochs. Visual scoring of the recordings were done following the current, widely used American Academy of Sleep Medicine scoring rules (AASM) [65] with requirements including slow waves to occupy at least 20% of a 30 second epoch in order to be classified as Stage 3. All scoring was performed using the SpiSOP tool (https://www.spisop.org; RRID: SCR_015673).

#### fMRI acquisition

For fMRI, functional images were acquired with ascending slice acquisition a T2*-weighted gradient-echo multiband echo-planar imaging sequence (Prisma 3T,Siemens,Erlangen, Germany; 66 axial slices; volume repetition time (TR), 1000 ms; echo time (TE), 34 ms; 60° flip angle; slice thickness, 2 mm; field of view (FOV) 210 mm; voxel size 2×2×2 mm). Anatomical images were acquired using a T1-weighted MP-RAGE sequence (192 sagittal slices; volume TR, 2300 ms; TE, 3.03 ms; 8° flip angle, slice thickness, 1 mm; FOV, 256 mm; voxel size 1×1×1 mm).

#### fMRI data processing

Image preprocessing and statistical analysis was performed using SPM8 software (www.fil.ion.ucl.ac.uk/spm; Wellcome Trust Centre for Neuroimaging, London, United Kingdom). All functional contrast images went through the standard preprocessing steps. The images were realigned, slice-time corrected, spatially normalized and transformed into a common space, as defined by the SPM2 Montreal Neurological Institute (MNI) T1 template. The preprocessed datasets were then analyzed using the general linear model and statistical parametric mapping [66]. The first five volumes from every dataset after preprocessing were discarded to remove non-steady state effects. For the statistical analysis, relevant contrast parameter images were generated for each subject and then subjected to a second-level full factorial analysis with nonsphericity correction for correlated repeated measures. For the first level analysis, individual contrast images for each subject were produced by comparing task dependent activation (hidden > cued) for each session separately, with six movement parameters as regressors of no interest. Since the time taken to complete the task was not uniform for all subjects, we performed the first level analysis on the activity in the first 30 seconds of every trial. Analyses was also done for the activity corresponding to the entire task length and is shown in the supplementary section. For the second level analyses, these contrast images were included in a full factorial model with between subject factors – Training (allo/ego) and Condition (sleep/wake) and within subject factor – session (training/test). In the whole brain search, all results were collected at p<0.005 uncorrected and then corrected at the cluster level to control for multiple comparisons (p<0.05 FWE-cluster).

For the functional connectivity analyses between training and test sessions, we chose the medial frontal cortex (coordinates 2 10 48) as the region of interest to capture network activity. The coordinates of the region were taken from the activity analysis where we observed an increase in activity in this region after sleep. This region has been previously shown to be a part of the executive control network and related to goal directed navigation [32]. A psycho-physiological interaction (PPI) analysis was performed. The psychological variable consisted of the activity of task blocks (hidden island) in the first 30 seconds of each training block convoluted to the hemodynamic response. The physiological factor was the time course of a spheroid volume of interest (VOI) located in the medial frontal cortex (2, 10, 48) with a 6 mm radius. The VOI time course was extracted for each individual and adjusted for head movement. With the PPI toolbox (SPM8) the interaction value (PPI) of both factors was calculated. The PPI, VOI time course and task timing was then included in a general linear model with the 6 movement parameters as regressors of no interest. For each participant, individual contrast images with the PPI activation were calculated. These contrast images were then included in a full factorial design model with the same factors as used for activity analysis. In the whole brain search, all results were collected at p<0.005 and then corrected at the cluster level to control for multiple comparisons (p<0.05 FEW-cluster). All data and analysis scripts will be available on the Donders Repository.

**Fig. S1.**
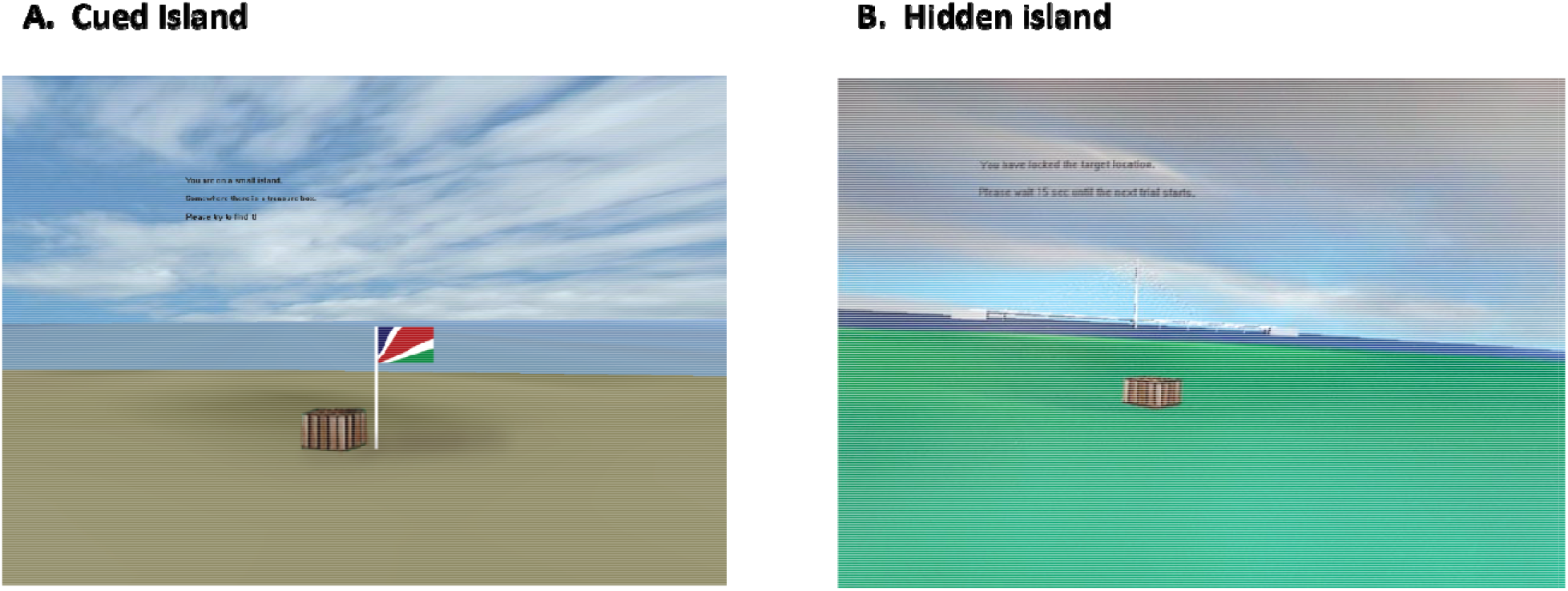
Virtual Watermaze. (A) The left panel displays a representation of the cued island. A plain brown island with no surrounding distal cues. The flag in picture is visible to the navigator from a distance and keeps changing it position each trial. (B) The right panel displays the target quadrant of the hidden island. A green island surrounded by four landmarks (distal cues) like the bridge as shown in the picture. The location of the box is fixed on this island and is located on an indentation in the virtual surface and only visible from close distance.

**Fig. S2.**
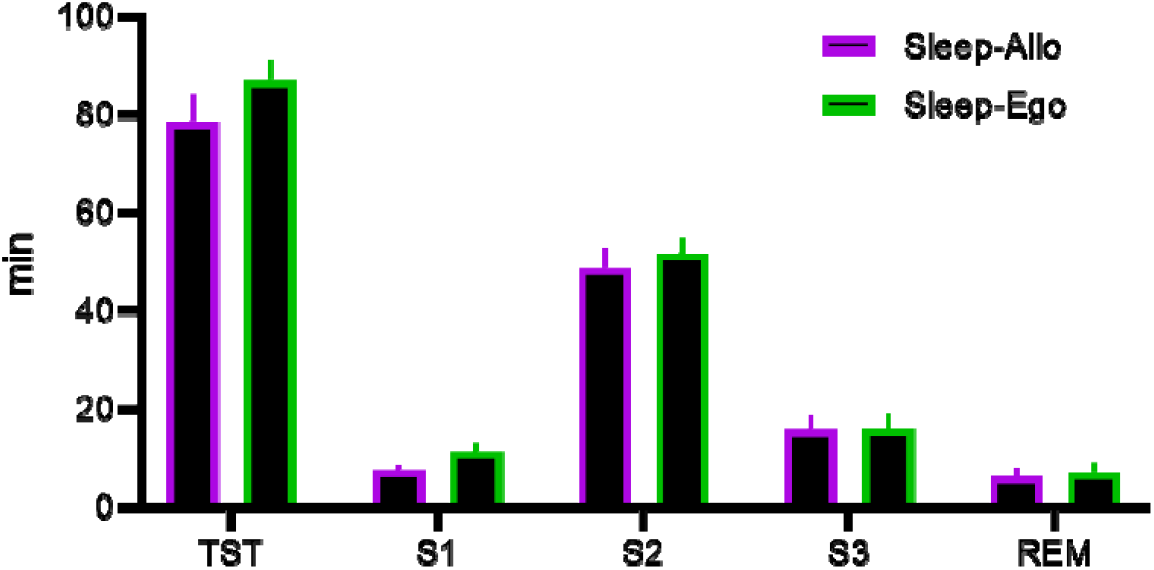
Human Sleep Stage Results. Shown is the amount of sleep during the nap for each training group. There was no significant difference between the two groups (multivariate analysis Allo vs Ego F_5,30_=0.9 p=0.52).

**Fig. S3.**
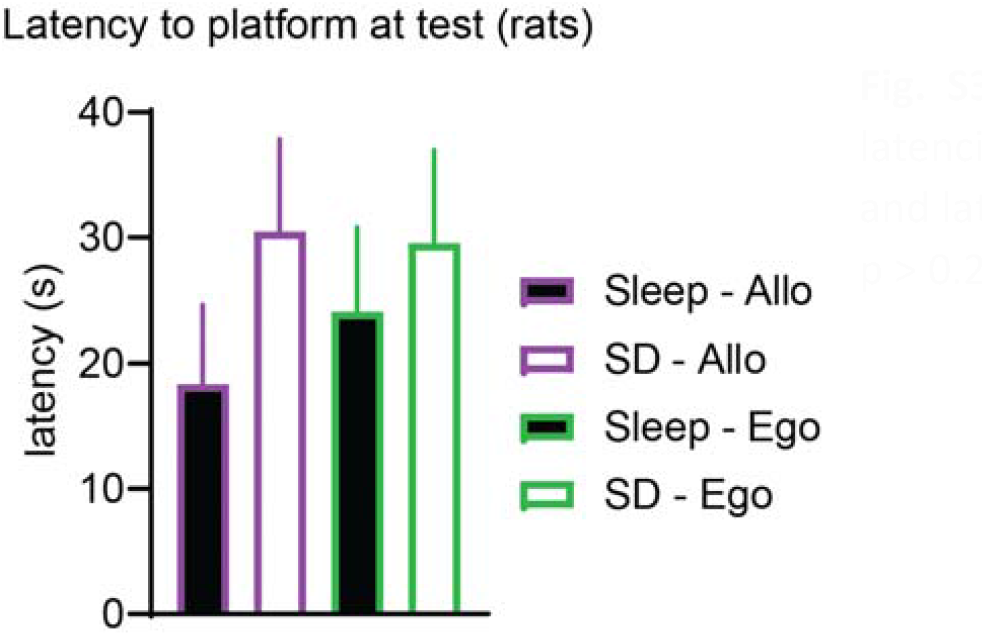
Latency to reach platform at test for rats. The latencies showed a similar pattern as dwell times in rats and latencies in humans, however it was not significant (all p > 0.2).

**Fig. S4.**
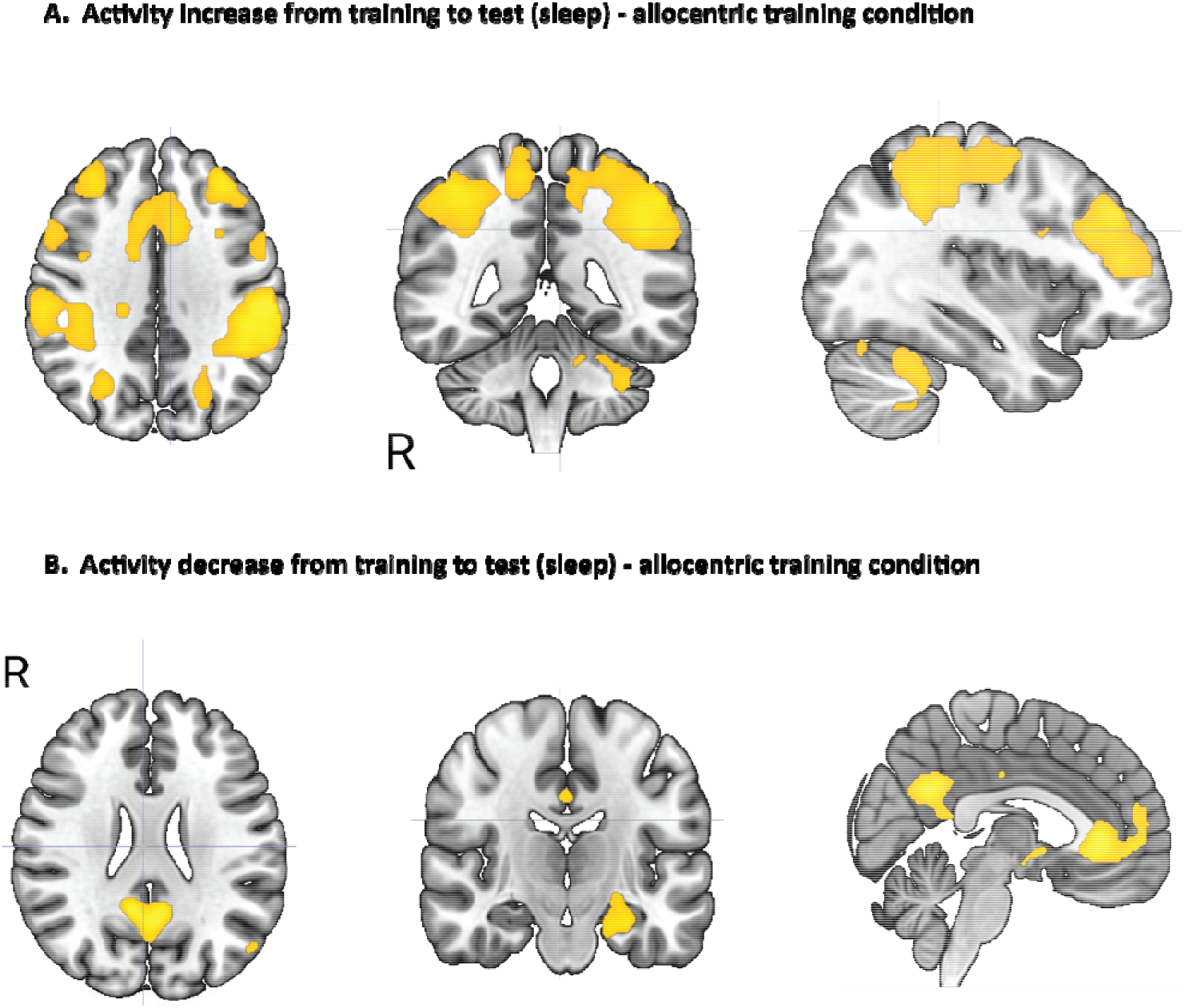
fMRI activity analysis in first 30 seconds of each training and test trial - Allocentric. Brain maps show the changes from the sleep group consisting of subjects who were trained under allocentric condition. (A) Regions with increased activation from training to test (test > training, p<0.05 FWE-cluster corrected). An increase in activity was observed in the medial and lateral frontal cortex, posterior parietal cortex. (B) Regions with decreased activation from training to test (test < training, p<0.05 FWE-cluster corrected). A decrease in activity was observed in the medial prefrontal cortex, precuneus and hippocampus. All results were collected at uncorrected p < 0.005 and then corrected on the cluster level to control for multiple comparisons with p<0.05 FWE

**Fig. S5.**
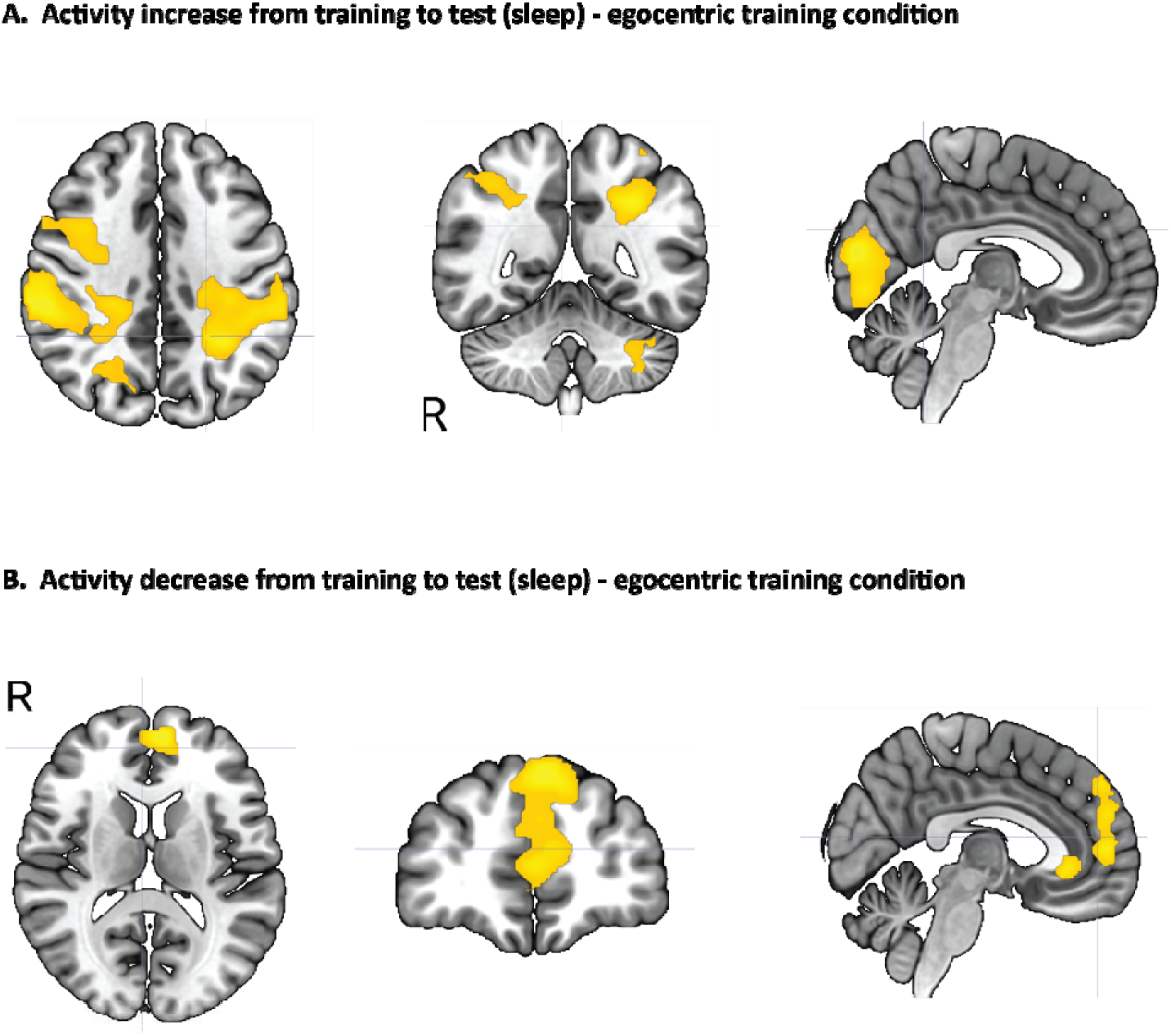
fMRI activity analysis in first 30 seconds of each training and test trial - Egocentric. Brain maps show the changes from the sleep group consisting of subjects who were trained under egocentri condition. (A) Regions with increased activation from training to test (test > training, p<0.05 FWE-cluster corrected). An increase in activity was observed in the anterior and posterior parietal cortex, a part of the visual cortex and cerebellum. (B) Regions with decreased activation from training to test (test < training, p<0.05 FWE-cluster corrected). A decrease in activity was observed in the medial prefrontal cortex and precuneus. All results were collected at uncorrected p < 0.005 and then corrected on the cluster level to control for multiple comparisons with p<0.05 FWE

**Fig. S6.**
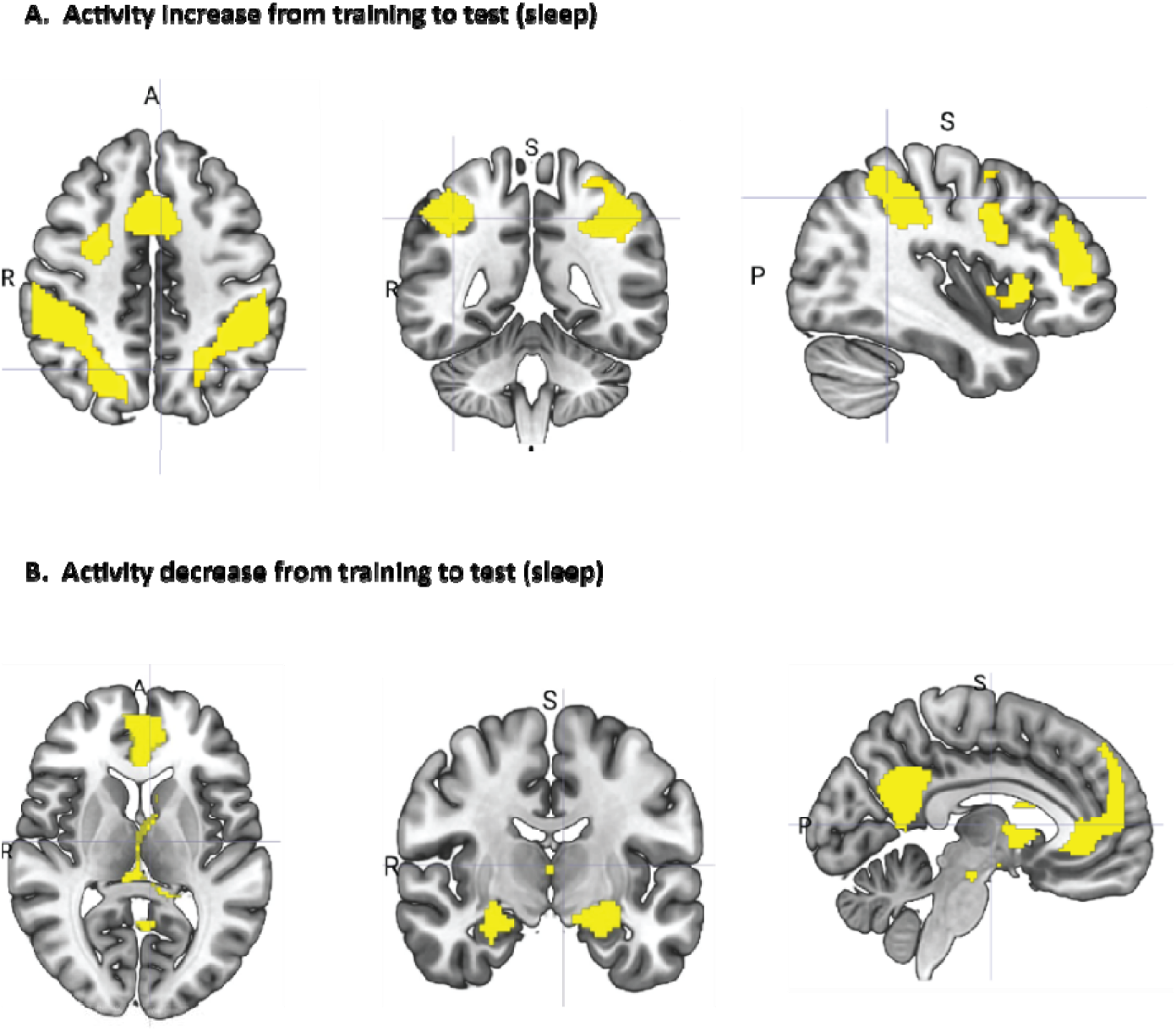
fMRI activity analysis for the entire length of each training and test trial. Brain maps show the changes from the sleep group consisting of subjects from both training conditions. (A) Regions with increased activation from training to test (test > training, p<0.05 FWE-cluster corrected). An increase in activity was observed in the medial and lateral frontal cortex, anterior and posterior parietal cortex. (B) Regions with decreased activation from training to test (test < training, p<0.05 FWE-cluster corrected). A decrease in activity was observed in the medial prefrontal cortex, precuneus and hippocampus. All results were collected at uncorrected p < 0.005 and then corrected on the cluster level to control for multiple comparisons with p<0.05 FWE

**Fig. S7.**
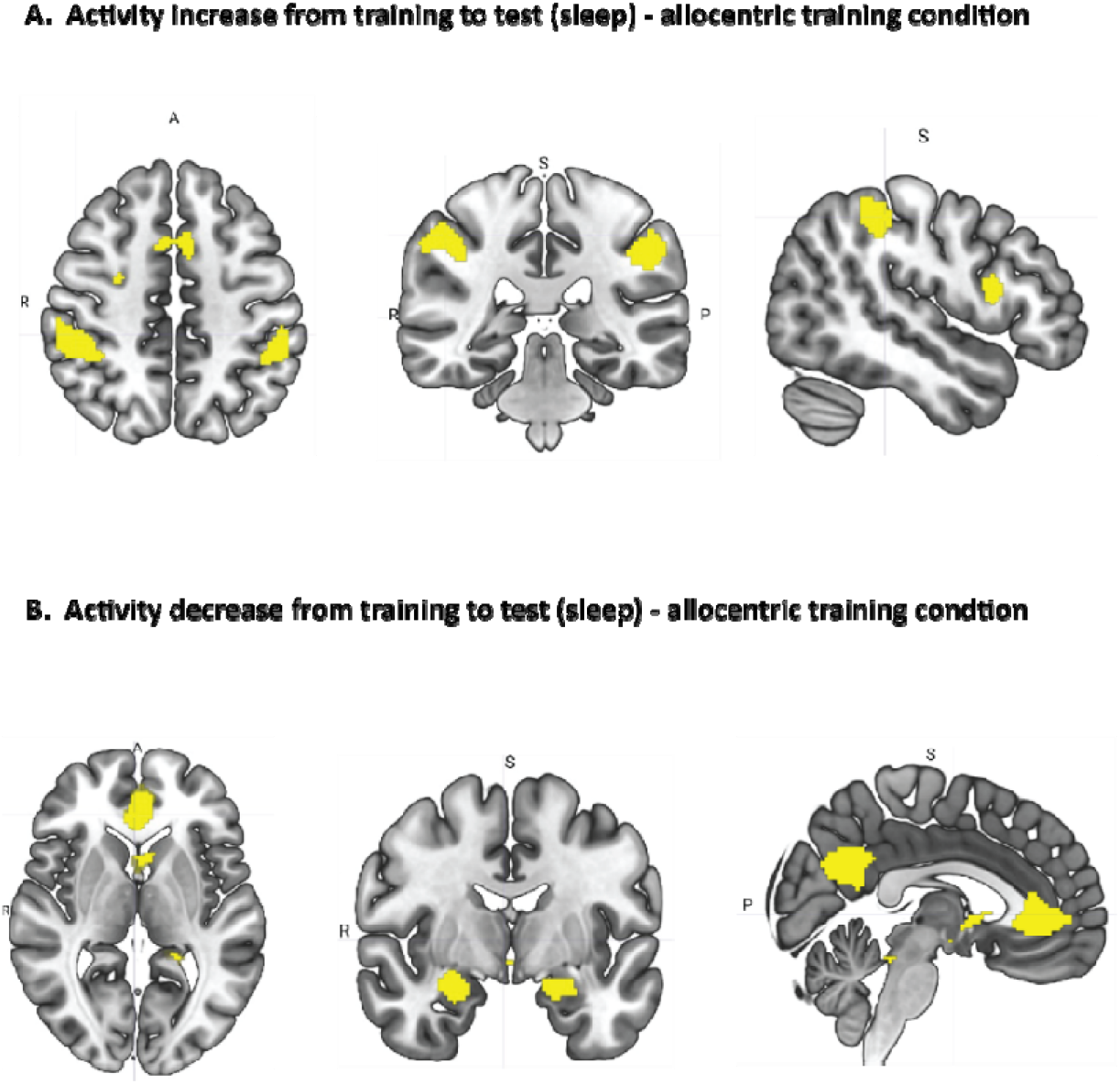
fMRI activity analysis for the entire length of each training and test trial - Allocentric. Brain maps show the changes from the sleep group consisting of subjects who were trained under allocentri condition. (A) Regions with increased activation from training to test (test > training, p<0.05 FWE-cluster corrected). An increase in activity was observed in the medial frontal cortex, posterior parietal cortex. (B) Regions with decreased activation from training to test (test < training, p<0.05 FWE-cluster corrected). A decrease in activity was observed in the medial prefrontal cortex, precuneus and hippocampus. All results were collected at uncorrected p < 0.005 and then corrected on the cluster level to control for multiple comparisons with p<0.05 FWE

**Fig. S8.**
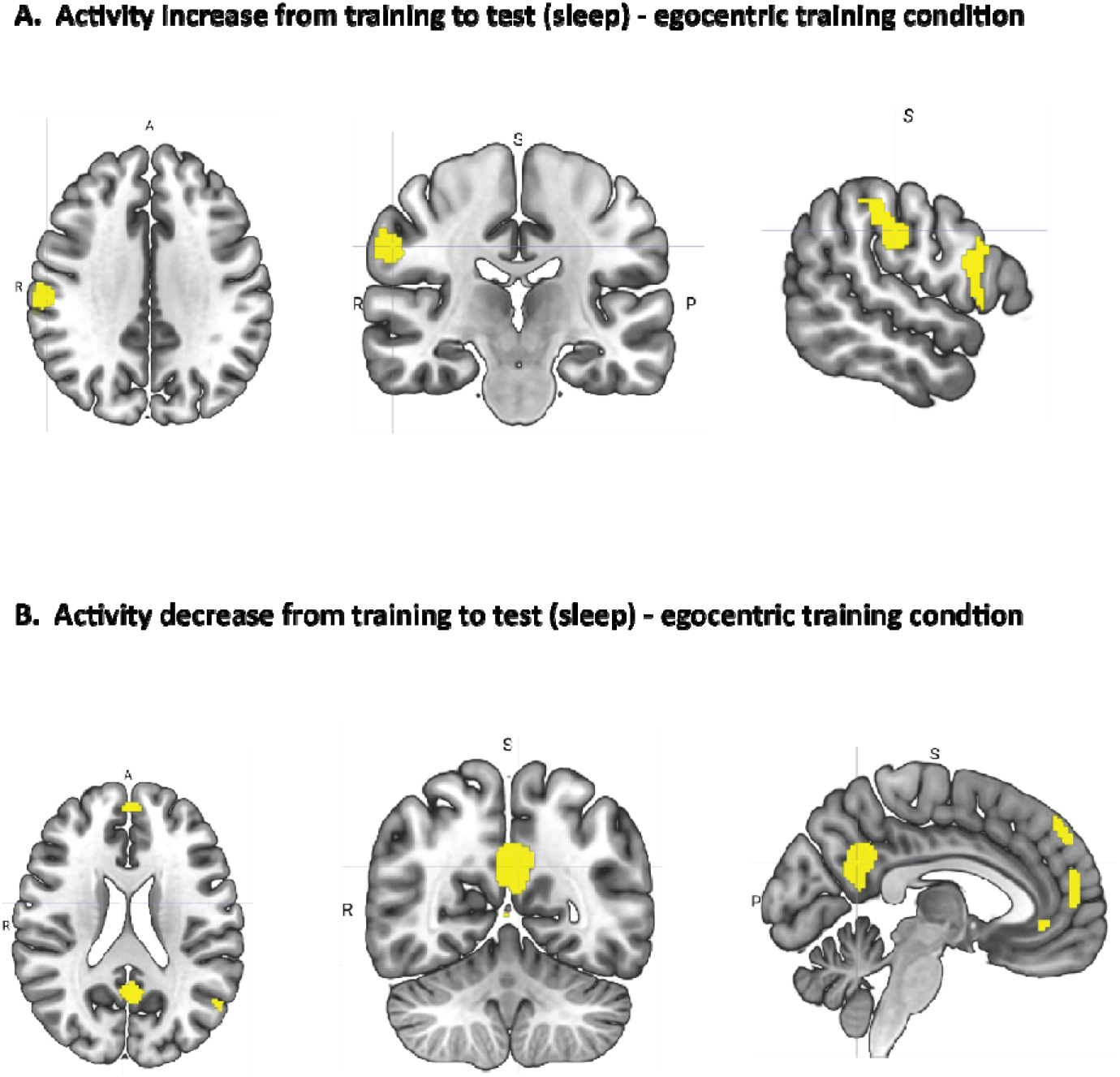
fMRI activity analysis for the entire length of each training and test trial - Egocentric. Brain maps show the changes from the sleep group consisting of subjects who were trained under egocentri condition. (A) Regions with increased activation from training to test (test > training, p<0.05 FWE-cluster corrected). An increase in activity was observed in the anterior and posterior parietal cortex. (B) Regions with decreased activation from training to test (test < training, p<0.05 FWE-cluster corrected). A decrease in activity was observed in the medial prefrontal cortex and precuneus. All results were collected at uncorrected p < 0.005 and then corrected on the cluster level to control for multiple comparisons with p<0.05 FWE

**Fig. S9.**
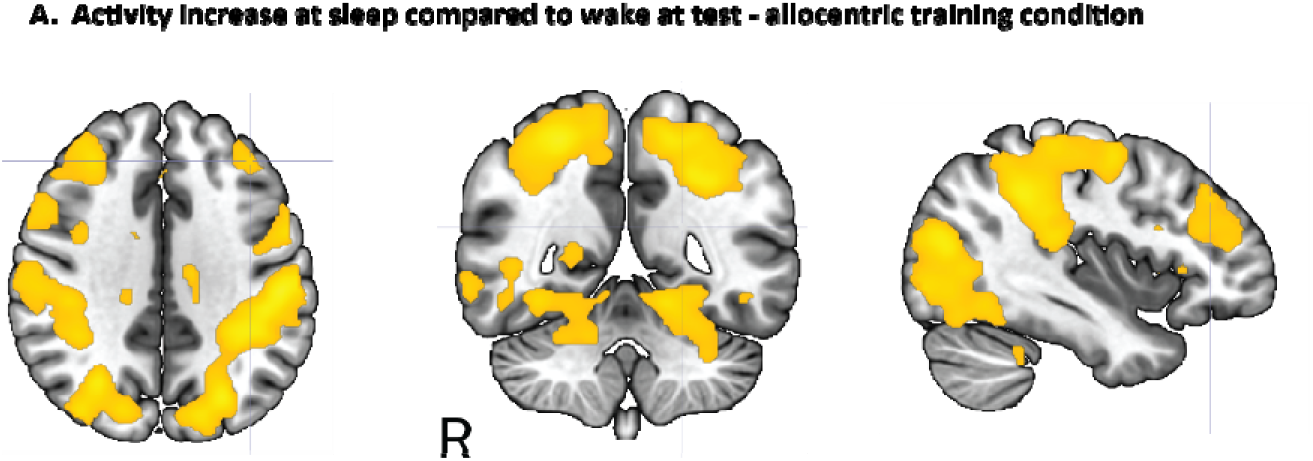
fMRI activity analysis for sleep vs wake groups at test trial – allocentric. Brain maps show changes in brain activity between subjects in sleep and wake groups for the test trial when trained under allocentric condition (sleep > wake, p<0.05 FWE-cluster corrected). An increase was observed in the medial and lateral frontal cortex, posterior parietal cortex. All results were collected at uncorrected p < 0.005 and then corrected on the cluster level to control for multiple comparisons with p < 0.05 FWE

**Fig. S10.**
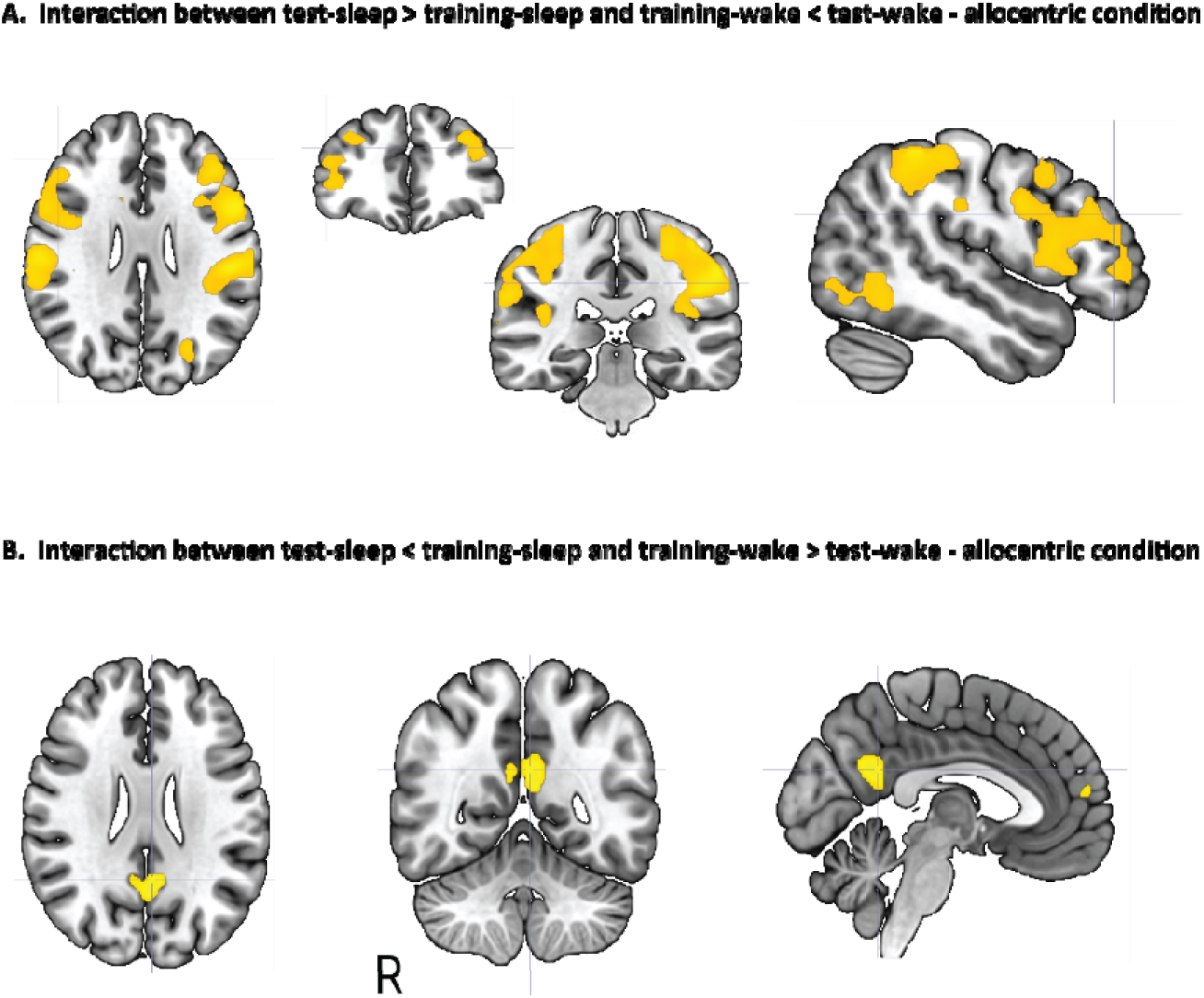
fMRI activity analysis for the interaction between sleep and wake groups at training and test – only allocentric. (A) Regions significant for test-S > training -S and training-W < test-W, p<0.05 FWE cluster corrected (B) The opposite contrast with test-S < training -S and training-W > test-W, results shown for uncorrected p<0.005. The clusters were not significant when whole brain cluster corrected.

**Fig. S11.**
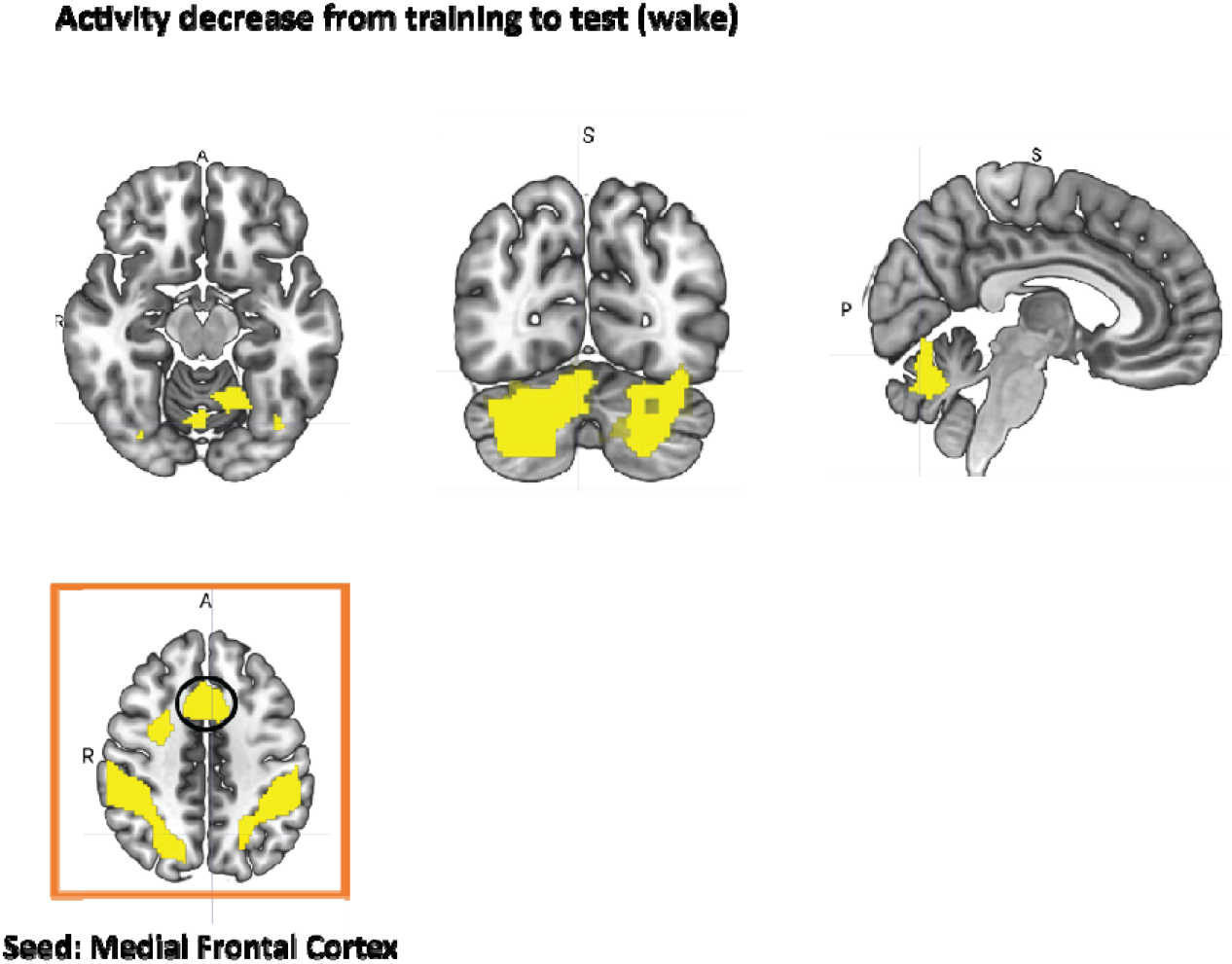
fMRI Connectivity Analysis (PPI) with Medial Frontal Cortex as ROI. Brain maps show changes from both wake groups (test<training, p<0.05 FWE-cluster corrected). The frontal medial cortex showed a significant decrease in connectivity with the cerebellum for both wake groups. All results were collected at uncorrected p < 0.005 and then corrected on the cluster level to control for multiple comparisons with p<0.05 FWE

**Table S1.** Demographic Data. Table S1 presents the background information of the human participants and their level of arousal before the first and second MRI sessions respectively.

|  | Sleep-Allocentric |  | Sleep-Egocentric |  | Wake-Allocentric |  | Wake-Egocentric |  |
| --- | --- | --- | --- | --- | --- | --- | --- | --- |
|  | M | SEM | M | SEM | M | SEM | M | SEM |
| Age (Years) | 23.06 | 0.74 | 21.9 | 0.56 | 24.8 | 0.81 | 23.31 | 0.75 |
| PSQI | 4.53 | 0.19 | 4.00 | 0.6 | 3.8 | 0.15 | 4.68 | 0.21 |
| SSS-1 | 2.08 | 0.18 | 2.00 | 0.68 | 2.46 | 0.23 | 2.15 | 0.19 |
| SSS-2 | 2.17 | 0.19 | 2.11 | 0.6 | 2.56 | 0.15 | 2.89 | 0.21 |

PSQI – Pittsburgh Sleep Quality Index ; SSS – Stanford Sleepiness Scale

**Table S2.**
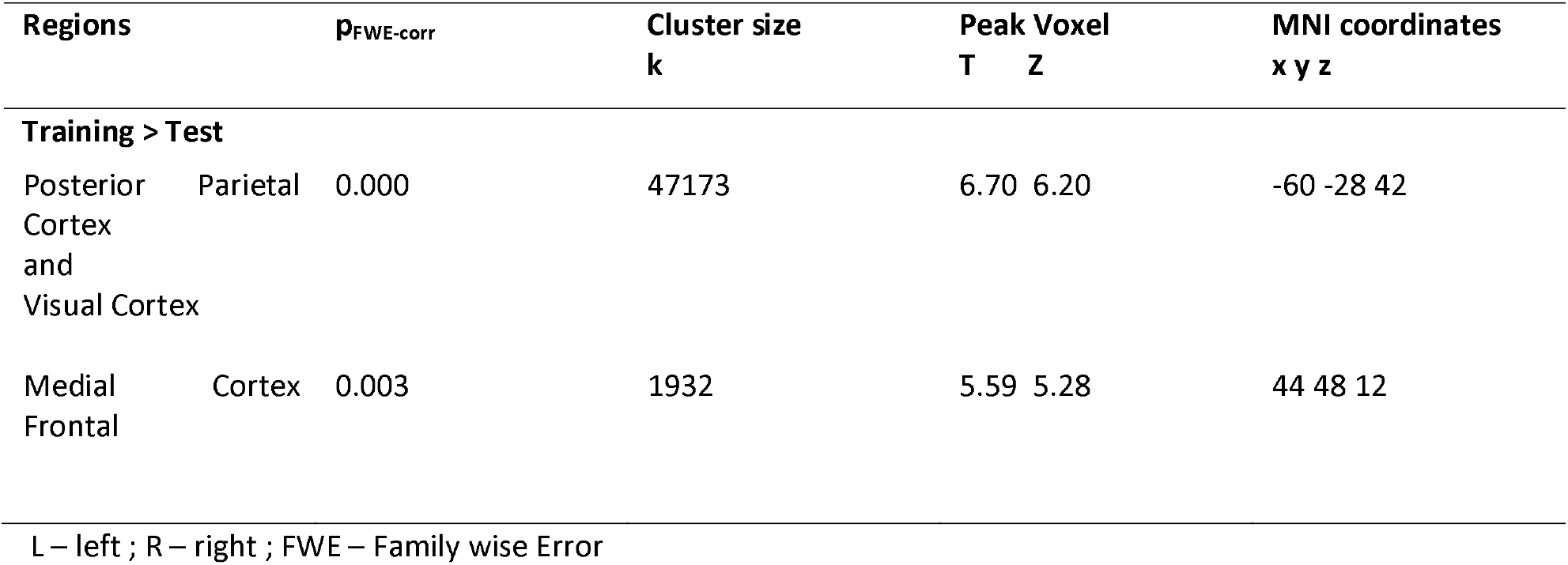
Activity Analysis – Increase in activity from training to test (sleep), p_FWE-cluster_<0.05, see Figure 3A

**Table S3.**
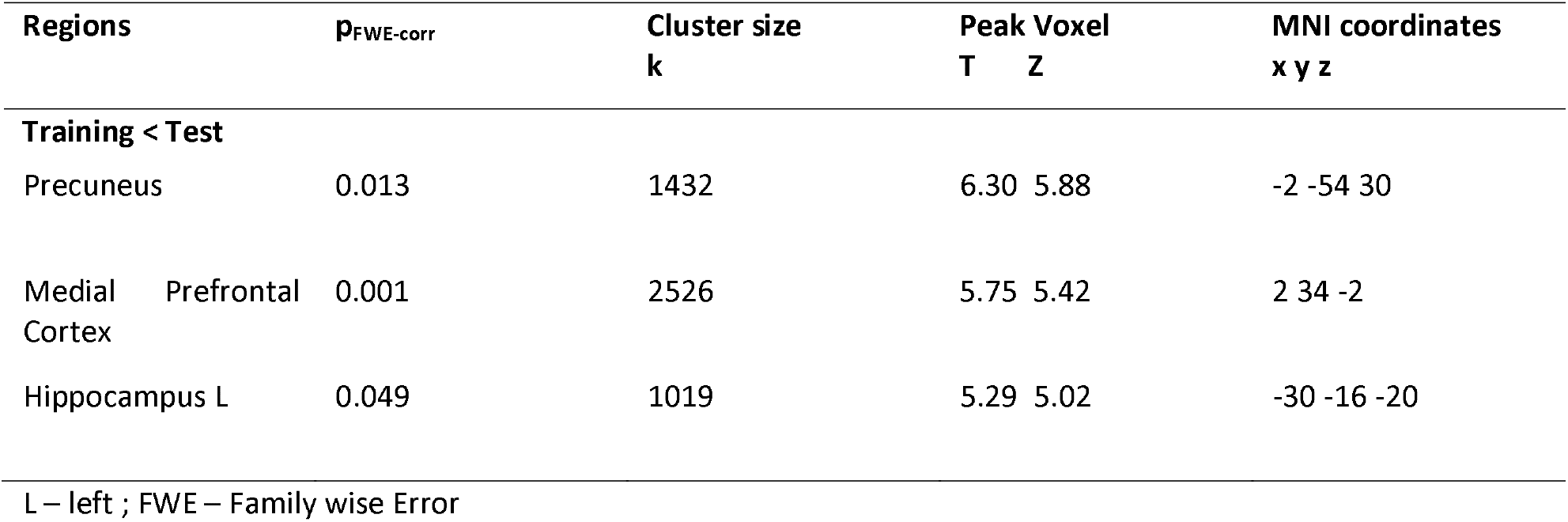
Activity Analysis – Decrease in activity from training to test (sleep), p_FWE-cluster_<0.05, see Figure 3B

**Table S4.**
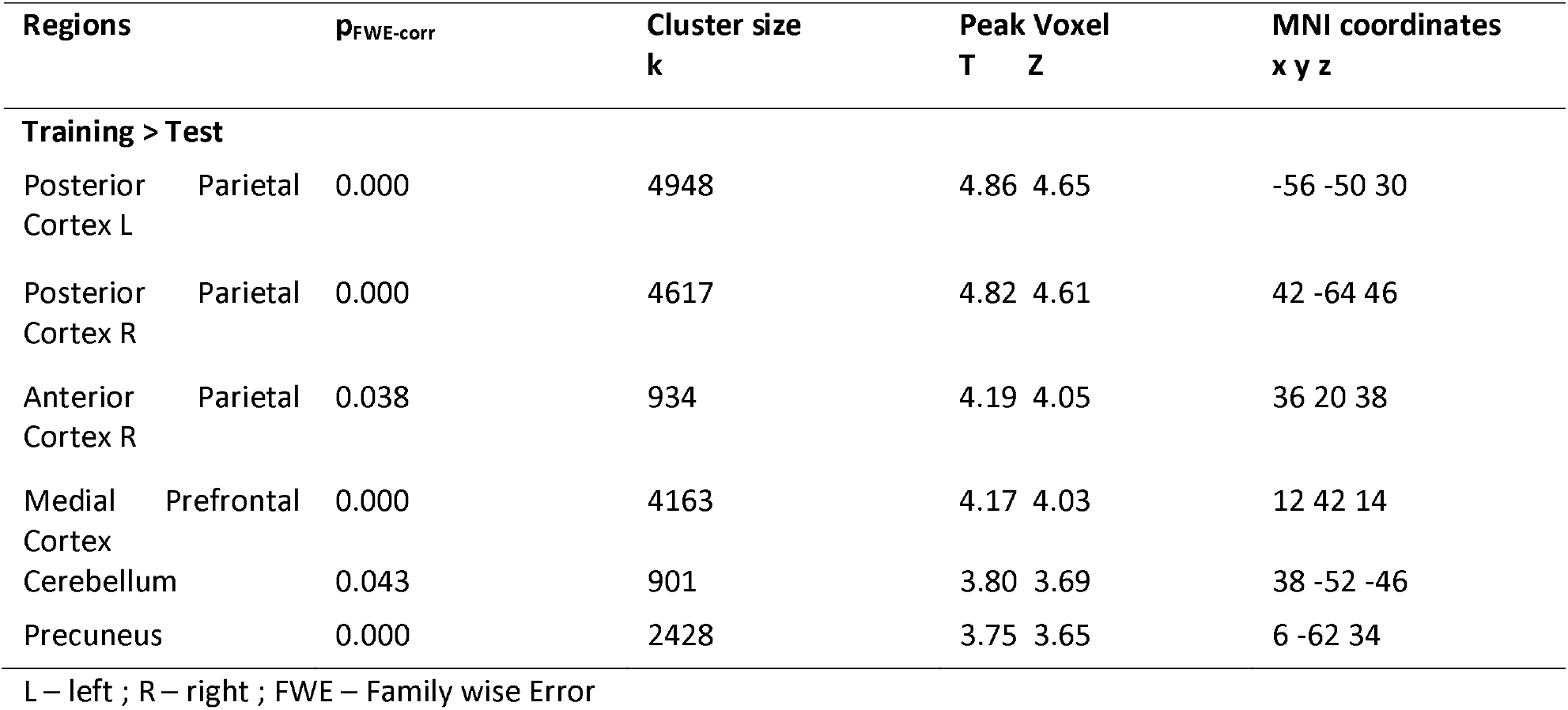
PPI Analysis – Connectivity decrease from Training to test (sleep), p_FWE-cluster_<0.05, see Figure 4

